# Diffusion-based Representation Integration for Foundation Models Improves Spatial Transcriptomics Analysis

**DOI:** 10.1101/2025.11.20.689624

**Authors:** Atishay Jain, Tuan M. Pham, David H. Laidlaw, Ying Ma, Ritambhara Singh

## Abstract

**Motivation:** We propose DRIFT, a framework that integrates spatial context into the input representations for foundation models by leveraging diffusion on spatial graphs derived from spatial transcriptomics (ST) data. ST captures gene expression profiles while preserving spatial context, enabling downstream analysis tasks such as cell-type annotation, clustering, and cross-sample alignment. However, due to its emerging nature, there are very few foundation models that can utilize ST data to generate embeddings generalizable across multiple tasks. Meanwhile, well-documented foundational models trained on large-scale single-cell gene expression (scRNA-seq) data have demonstrated generalizable performance across scRNA-seq assays, tissues, and tasks; however, they do not leverage the spatial information in ST data. We use heat kernel diffusion to propagate embeddings across spatial neighborhoods, incorporating the local neighborhood context of the ST data while preserving the transcriptomic representations learned by state-of-the-art single-cell foundation models.

**Results:** We systematically benchmark five foundational models (both scRNA-seq and ST-based) across key ST tasks such as annotation, alignment, and clustering, ensuring a comprehensive evaluation of our proposed framework. Our results show that DRIFT significantly improves the performance of existing foundational models on ST data over specialized state-of-the-art methods. Overall, DRIFT is an effective, accessible, and generalizable framework that bridges the gap toward universal models for modeling spatial transcriptomics.

**Availability and Implementation:** Code and data available at https://github.com/rsinghlab/DRIFT.

**Supplementary information:** Supplementary notes are provided with the manuscript.

## Introduction

Spatial transcriptomics (ST) technologies enable measurement of gene expression while preserving tissue architecture, offering a direct view of how cellular identity and spatial organization together shape biological function. By jointly capturing transcriptomic and spatial information, ST datasets reveal the local microenvironment of gene activity and support downstream analyses such as spatially variable gene detection [8, 1, 46, 23, 54], clustering [47, 50, 28, 13, 52], cell–cell interaction inference [44, 5, 22, 16], and cross-section alignment [48, 27]. These capabilities have made ST an essential approach in developmental biology, pathology, and tissue-level systems analysis. However, the rapidly growing complexity and heterogeneity of ST data necessitate generalizing and automating the downstream tasks across a wide range of tissues, species, and technologies. Furthermore, ST measurements are typically sparse, noisy, and less throughput-efficient than single-cell RNA sequencing (scRNA-seq). This highlights the need for robust computational models that can extract biologically meaningful and spatially aware representations that generalize to diverse downstream ST analyses.

Recently, to address these challenges, ST foundation models have emerged, marking a growing effort to bring foundation modeling into the ST domain. Early examples include Loki [11], Nicheformer [34], SToFM [53], and STFormer [24]. Loki aligns histology images and spatial gene-expression profiles through cross-modal contrastive learning; Nicheformer applies a transformer trained on ST data, where attention mechanisms capture neighborhood-level dependencies, and SToFM and STFormer introduce spatial attention and positional encoding schemes to jointly model biological and spatial information. However, they rely heavily on large paired multimodal datasets [11] or cell-type priors derived from deconvolution [24], which are often unavailable for many ST platforms. Moreover, some models do not explicitly model spatial relationships, limiting their ability to capture cell neighborhoods [11, 34]. Additionally, training these architectures from scratch is computationally expensive, often requiring substantial computing resources. Finally, existing ST foundation models are trained on relatively homogeneous and small ST datasets. As a result, these models typically require dataset-specific retraining, limiting their utility as universal foundation models for ST datasets.

On the other hand, scRNA-seq foundation models are trained on millions of scRNA-seq profiles from diverse tissues and species. Models such as Geneformer [37, 10], scGPT [12], and scFoundation [17] learn gene–gene dependencies using transformer-based or masked language-model architectures. The embeddings generated by these models capture regulatory patterns, providing general-purpose representations that can transfer effectively across datasets and biological contexts. Their success demonstrates that large pretrained models can generalize across data sources and experimental conditions with limited fine-tuning, offering a promising foundation for diverse multi-tissue and cross-species analyses [43]. However, since these models are trained solely on scRNA-seq data, they lack the spatial context critical for modeling ST data.

We introduce **D**iffusion-based **R**epresentation **I**ntegration for **F**oundation models in spatial **T**ranscriptomics (**DRIFT**). DRIFT constructs a spatial adjacency graph among tissue spots (or cells) and applies a heat-kernel diffusion process that propagates gene-expression signals across local neighborhoods while preserving tissue boundaries. This smoothing produces spatially coherent and denoised gene expression representations that can be directly fed into any pretrained foundation model without retraining, making our approach much more computationally scalable and accessible. Foundation models that do not explicitly model neighborhood information benefit from both spatial context integration and denoising, while methods that do so leverage DRIFT’s denoised input.

We demonstrate that DRIFT generalizes easily across three critical ST tasks: cell-type annotation, cross-section alignment, and clustering. A single DRIFT framework outperforms state-of-the-art methods specialized for the different ST tasks, thereby setting a new norm for ST downstream analysis. Furthermore, the DRIFT diffusion leverages existing pretrained scRNA-seq foundation models, achieving improved performance compared to baseline ST foundation models. Therefore, our results establish that the diffusion operator at the core of DRIFT provides a simple, efficient, and general way to couple pretrained scRNA-seq foundation models with ST data by incorporating a spatially aware representation. For ST foundation models that may embed spatial context, diffusion-based denoising still improves performance. DRIFT’s generalizability across technologies, tissues, and models underscores its robustness for the evolving ST field.

## Method

As illustrated in Figure 1, we introduce a graph diffusion-based framework, DRIFT, to incorporate neighborhood information into the gene expression profiles. To perform this integration, we construct a spatial graph and apply heat kernel diffusion over it. Our method operates on the premise that spatially proximal cells/spots likely share cell-type identity, allowing us to denoise the data by propagating signal across local neighborhoods [45, 36, 33]. The diffused gene expression matrices obtained from DRIFT are used as inputs to pretrained foundation models, which are then evaluated for multiple important ST analysis tasks over a variety of ST datasets.

**Fig. 1.**
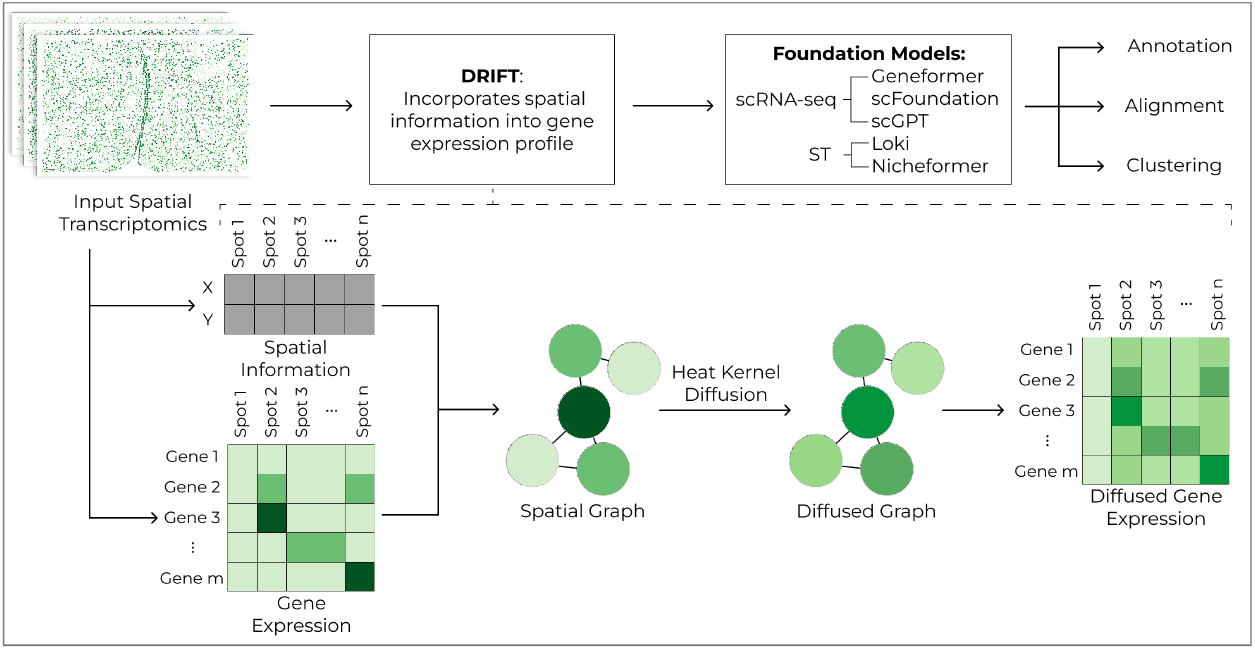
An illustration of the DRIFT Framework. Gene expression profiles of individual spots do not include any spatial context. Therefore, we create a spatial graph in which each node represents a cell/spot, and each edge represents the spatial proximity between the corresponding nodes. We then apply heat kernel graph diffusion on the constructed graph and use the output diffused gene expression profiles for each cell/spot. Through diffusion, each node incorporates information about its neighbors. The diffused gene expression values are fed into foundation models instead of the original gene expression for downstream tasks.

### DRIFT Approach

First, we represent the ST data as a graph *G* = (*V, E*), where each node *v*_*i*_ ∈ *V* corresponds to a single tissue location in the ST dataset (i.e., a cell or spot). Edges (*v*_*i*_, *v*_*j*_) ∈ *E* connect spatially proximal nodes based on Euclidean distance. This graph construction strategy is applied uniformly across both cell-resolution and spot-resolution datasets. Following an empirical evaluation of construction strategies (detailed in Supplementary Section S1), we implemented a *k*-nearest neighbor graph with *k* = 3, which yielded the most robust performance across datasets irrespective of ST platform resolution. The edges of this graph are represented by the adjacency matrix *A* ∈ ℝ^*n*×*n*^, where *A*_*ij*_ = 1 if nodes *i* and *j* are spatial neighbors and *A*_*ij*_ = 0 otherwise.

We then calculate the degree matrix *D*, which is a diagonal matrix defined as

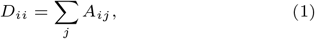

representing the degree of each node *i*. Using these, we compute the graph Laplacian as

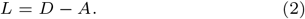

The Laplacian encodes the structure of *G* and serves as a discrete analogue of the continuous Laplace operator, capturing how information flows between connected nodes.

Heat kernel diffusion models the propagation of information (or “heat”) across a graph. It is defined by the heat equation on graphs:

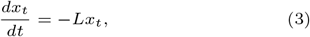

Solving this differential equation yields the heat kernel solution:

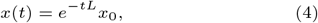

where, *x*_0_ represents the original (non-diffused) gene expression matrix and *x*_*t*_ denotes the diffused gene expression matrix at time *t. t* is therefore a hyperparameter that controls the strength of diffusion. Small *t* values (representing a shorter time for heat diffusion) lead to weaker diffusion and preserve local structure. Meanwhile, large *t* values result in stronger smoothing across larger spatial distances.

#### Relation to Spectral Graph Theory

Since *G* is an undirected graph, its Laplacian *L* is a symmetric real matrix and is therefore diagonalizable. By the graph spectral theorem, it can be decomposed as

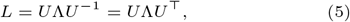

where, *U* is the orthogonal matrix of eigenvectors and Λ is the diagonal matrix of the corresponding eigenvalues. Substituting equation 5 into the heat kernel (equation 4) gives:

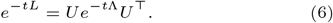

This expression reveals that diffusion corresponds to exponentially decaying the eigenvalues of the Laplacian, which is equivalent to a soft low-pass filter on the graph spectrum. Effectively, it smoothens gene expression based on the graph structure [49], incorporating the cell’s neighborhood information. This idea has been previously explored for scRNA-seq [40] and 3D genome organization data [39, 18], further demonstrating that graph diffusion techniques can effectively denoise data, smooth signals across neighboring nodes, and capture spatial relationships. The resulting diffused gene expression matrix *x*(*t*) thus integrates the spatial context of cells directly with the gene expression matrix, making it compatible with existing scRNA-seq foundation models that only accept gene expression values as inputs. Therefore, our choice of heat kernel diffusion is appropriate for our ST data modeling task. To further justify our choice of the heat kernel for graph diffusion, we show that it outperforms other diffusion and denoising methods overall (Supplementary Section S2).

Supplementary Section S3 validates the importance of spatial context via graph shuffling and neighborhood prediction benchmarks, confirming that DRIFT successfully incorporates this context into the input gene expression profiles. Next, we input the diffused gene expression matrix into the foundation models.

### Foundation Models

We apply DRIFT to a variety of scRNA-seq and ST foundation models to illustrate its ability to leverage the performance of existing models. We first demonstrate that DRIFT incorporates neighborhood information by pairing it with scRNA-seq foundation models. We further show its denoising property by enhancing the performance of ST foundation models.

#### Geneformer

Geneformer is a scRNA-seq foundation model that has three models. V1 is trained on 30 million cells [37], while V2 is trained on 104 million cells [10]. Furthermore, V2 has two model architecture configurations: one with 104 million parameters and another with 316 million parameters. In our Supplementary Section S4, we demonstrate that V2, with 316 million parameters, consistently outperforms other Geneformer configurations. Therefore, for any further experiments, we select V2 with 316 million parameters as the Geneformer foundation model.

#### scFoundation

scFoundation is a large-scale foundation model pretrained over 50 million single-cell transcriptomic profiles with an underlying encoder–decoder architecture.

#### scGPT

scGPT is a scRNA-seq foundation model trained on over 33 million cells, simultaneously learning cell and gene embeddings. It utilizes a specially designed attention mask and generative training to jointly optimize both embeddings.

#### Loki

Loki is a visual-omics foundation model that learns joint representations between histology images and transcriptomic profiles using contrastive learning based on OpenClip [21]. It is trained on 2.2 million paired image-transcriptomic patches. The framework does not explicitly train spatial information.

#### Nicheformer

Nicheformer is a transformer-based foundation model trained on 110 million single-cells and ST spots. By training on both scRNA-seq and ST, Nicheformer learns the characteristics of different dataset types. Nicheformer also does not explicitly use spatial information during training.

We provide additional technical context in Supplementary Section S5, including a brief explanation of how the foundation models handle varying gene sets across different ST platforms. The embeddings from these foundation models are then used in three important downstream ST tasks, as described below.

### Spatial Transcriptomic Tasks and Specialized Methods

To demonstrate the advantage of DRIFT, we measure its performance improvement over baseline foundation models across multiple ST tasks. These tasks are crucial for analyzing ST data and also facilitate cell-atlas construction, understanding disease mechanisms, and biomedical discovery [38, 42]. We also compare the DRIFT-incorporated foundation models to well-documented state-of-the-art specialized methods for each task.

#### Cell-type annotation

The aim of cell-type annotation is to identify cell types in a supervised manner, given a reference dataset. This is critical for understanding the heterogeneity of tissues and related biological processes [14]. Annotation is formulated as a few-shot learning task, where we attach a classification head (or probe) to a pretrained foundation model, and only train the weights of the probe to identify the cell types. We compare against STELLAR [4], a widely adopted deep-learning framework that has demonstrated superior annotation performance. STELLAR generates spatial graphs to represent ST data and uses a graph neural network (GNN) to learn embeddings of spots in an annotated reference ST dataset. Then it annotates cells in the unannotated ST dataset using the embeddings it generates. We assess the annotation performance through the accuracy and F1-score metrics.

#### Slice alignment

ST slice alignment is essential for reconstructing coherent anatomical structures from adjacent ST sections, as these slices represent slightly different structures and are subject to technical rotation, stretching, or deformation. Evaluating alignment accuracy, therefore, provides a rigorous test of how well DRIFT-incorporated embeddings preserve spatial geometry and enable meaningful cross-slice correspondence. We treat this as a zero-shot setting, applying Coherent Point Drift (CPD) directly to the original and DRIFT-incorporated embeddings, without any task-specific fine-tuning, across three alignment contexts with varying levels of noise and distortion. Baseline task-specific methods are PASTE [48] and PASTE2 [27]. PASTE uses Fused Gromov–Wasserstein (FGW) alignment, returning probabilistic alignment, while PASTE2 extends this to partial FGW alignment to handle partial overlaps. They are best suited for spatial alignments for homogeneous datasets (i.e. across continuous slices) [20].

We evaluate alignment performance using DRIFT-incorporated embeddings across both simulated and real-world datasets. In the simulation experiment, we construct each cell/spot’s total count using a Negative Binomial fit to the real library-size mean and variance, then generate per-gene counts by drawing from a multinomial whose probabilities are the real gene proportions with a pseudocount *δ* added.

For simulated data, we evaluate alignment quality using translation error together with PCC and Kendall’s *τ*, since ground-truth spatial coordinates are available. For real-world datasets, where true spatial coordinates are unknown, we instead assess alignment through LTARI, PCC, and Kendall’s *τ*. We average these metrics into a single alignment score reflecting both global and local alignment quality.

#### Translation Error

The translation error, defined as ∥*t*_*pred*_ − *t*_*true*_∥_2_ measures the Euclidean discrepancy between the predicted translation vector and the ground-truth shift required for correct spatial alignment. A lower translation error indicates a more accurate alignment compared to the ground truth. Translation error is computed only when ground-truth spatial coordinates are available (simulated data).

#### Pearson Correlation

For each aligned pair, we compute the Pearson correlation coefficient (PCC) as follows:

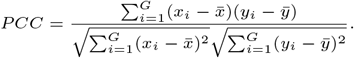

PCC captures the global expression pattern and linear correspondence between aligned slices. We report the median PCC across all aligned pairs, with higher values indicating stronger transcriptomic consistency between matched cells.

#### Kendall’s *τ*

We quantify rank-based concordance between aligned cells using Kendall’s 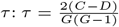, where *C* and *D* are counts of concordant and discordant gene pairs, and a higher *τ* indicates stronger rank similarity. We report the median *τ*, with higher values suggesting biologically coherent correspondence between matched cells.

#### Label Transfer Adjusted Rand Index (LTARI)

We compute LTARI as the adjusted Rand index between true labels **l** and alignment-transferred labels **Î** when labels are available. The mathematical formulation is LTARI = ARI(**l, Î**). A higher LTARI score indicates a more coherent local structure and cell-type organization across alignments.

#### Alignment Score

We compute an overall aggregate alignment score by averaging normalized PCC, Kendall’s *τ*, and LTARI values. By integrating both global and local metrics into a single score, the alignment score provides a quantitative analysis of overall alignment performance.

#### Clustering

Unsupervised clustering is the task of grouping cells without any prior information, which is an essential step for identifying marker genes [19, 25], and understanding disease pathology [9, 6]. Since it does not explicitly require any labels for training, it is a zero-shot task in which embeddings obtained from the foundation models are directly fed into unsupervised clustering algorithms, such as mclust [35] (a finite normal mixture modeling-based clustering algorithm), to obtain the final clusters.

To benchmark clustering performance, we compared DRIFT against GraphST [28], which has been identified as a top-performing method in recent independent evaluations [47, 20]. GraphST employs a GNN autoencoder architecture and contrastive learning techniques to learn embeddings that are subsequently utilized in unsupervised algorithms. We evaluate the methods using the adjusted Rand index (ARI). ARI is a metric commonly used to assess unsupervised clustering performance [28, 47, 20] because it accounts for the risk of accidental cluster agreement.

### Datasets

To demonstrate DRIFT’s generalizability, we benchmark it across eight datasets from various ST technologies and tissues of humans and mice. The ST technology, species, tissue, and number of slices for all the datasets are summarized in Table 1.

**Table 1.**
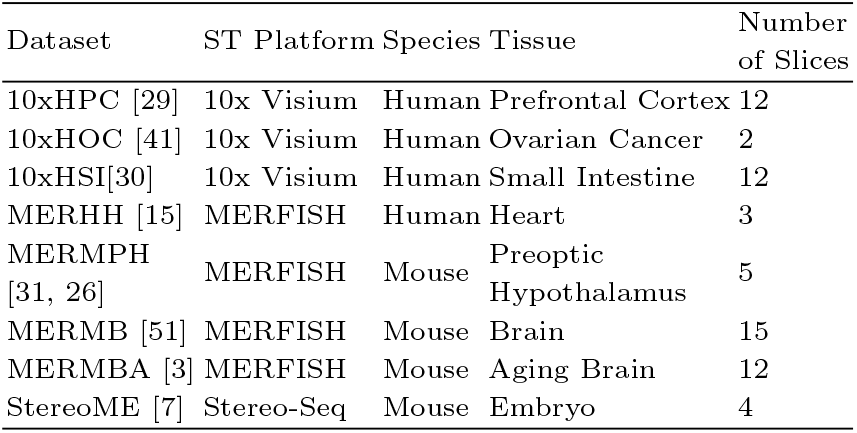
A summary of all the datasets. The table lists the ST technology, species, tissue, and number of slices in all the datasets.

### Hyperparameter Tuning

For every task, we tuned the *t* value (representing the strength of diffusion) in the heat kernel diffusion equation. The values assigned to *t* during hyperparameter tuning were 0.00001, 0.0001, 0.001, 0.01, 0.1, 1, 2, 5, 10, 100. We observed that *t* values from 1 to 10 performed similarly well overall. For alignment, we recommend selecting the *t* that yields the highest PCC (forfeiting the need for labels). However, for annotation and clustering, having a ground truth to tune *t* may be unfeasible. In such a case, we suggest a default value of *t* = 5 due to its high performance across tasks, datasets, and foundation models. Results for the hyperparameter tuning of *t* are summarized in Supplementary Section S6.

Any further hyperparameter tuning necessary for both foundation models and specialized methods during few-shot learning tasks was performed using Optuna [2] with 10 trials. We also tuned PASTE and PASTE2 for the zero-shot alignment task. The hyperparameters and their value ranges are listed in Supplementary Sections S7 and S8.

## Results

### DRIFT demonstrates state-of-the-art cell-type annotation performance

We first tested DRIFT’s performance on the cell-type annotation task. We formulated it as a few-shot problem, with training performed on just one slice. For all foundation models, we attached a three-layer multilayer perceptron (MLP) classification head to predict cell-type labels, while freezing all their encoder weights. The MLP was trained using the cross-entropy loss.

In this experiment, we employed datasets 10xHPC (10x human prefrontal cortex), MERHH (MERFISH human heart), MERMPH (MERFISH mouse preoptic hypothalamus), MERMB (MERFISH mouse brain), and MERMBA (MERFISH mouse brain) since they contain expert annotations necessary for evaluating the methods. For spot-resolution platforms, annotations represent the dominant cell type within the spot, as determined by the provided expert pathology annotations. The five foundation model classification heads and one specialized method (STELLAR) were trained and evaluated independently on each dataset. To recreate the realistic scenario of limited annotated data, we used only one annotated slice per dataset to train and hyperparameter-tune the methods, reserving the remaining slices for testing.

Figure 2 displays DRIFT’s F1-score performance improvement across all methods and datasets, demonstrating that the heat kernel’s spatial context integration and denoising properties are beneficial for the annotation task. We first compared the original scRNA-seq foundation models to the corresponding DRIFT-incorporated foundation models. Compared to the original foundation models, DRIFT shows average improvements of 19.81% to 32.5% in accuracy and 30.75% to 40.57% in F1-score. Dataset-specific accuracy results also show the same performance trends and are plotted in Supplementary Section S9.

**Fig. 2.**
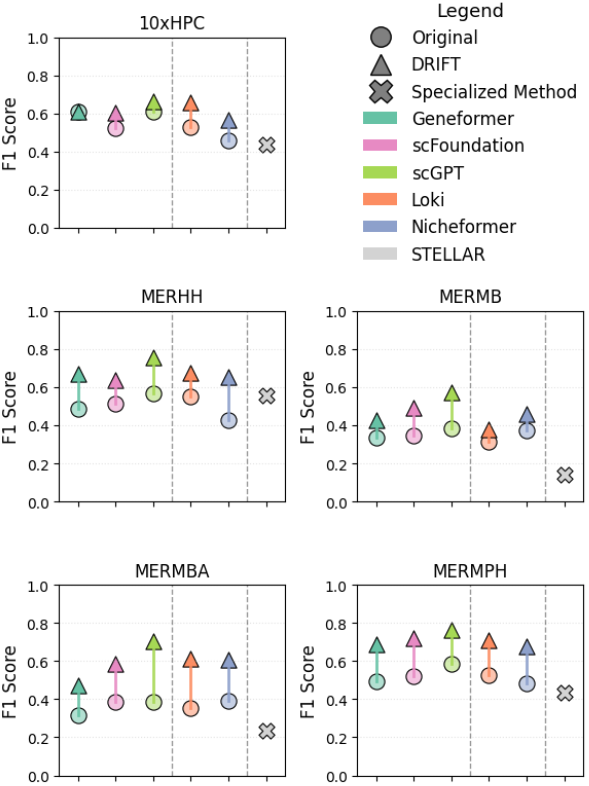
Cell-type Annotation Results. A quantitative comparison across multiple datasets. The circles represent the performance of the original foundation models, the triangles represent foundation models with DRIFT-enhanced inputs, and the X represents the specialized method STELLAR. The colors indicate which column corresponds to each foundation model (the order of models in the legend matches column placement). This figure shows that DRIFT improves the F1 score of foundation models on the annotation task. Furthermore, DRIFT-incorporated foundation models, specifically scGPT show state-of-the-art performance, surpassing even the specialized model - STELLAR.

In particular, scGPT with DRIFT shows the highest performance over all datasets. Furthermore, it consistently outperforms the specialized method, STELLAR. scGPT with DRIFT shows, on average, a 73.32% improvement in accuracy, and a 131.89% improvement in F1-score. scGPT with DRIFT also outperforms the original ST-based Loki and Nicheformer models. When compared to Loki, DRIFT-incorporated scGPT shows an average accuracy improvement of 48.46% and an average F1-score improvement of 57.96%. Similarly, when compared to Nicheformer, it shows an average accuracy improvement of 50.86% and an average F1-score improvement of 62.65%. These results demonstrate its state-of-the-art performance for ST annotation.

DRIFT also shows an improved performance for both the ST foundation models. Compared with the original Loki model, DRIFT-incorporated Loki shows an average accuracy improvement of 32.01% and an average F1-score improvement of 35.28%. Similarly, for Nicheformer, DRIFT shows an average accuracy improvement of 30.67% and an average F1-score improvement of 38.88%. These results illustrate the effectiveness of DRIFT in enhancing the ST foundation models.

### DRIFT improves ST alignment across different contexts

For alignment tasks, we set up three experiments on one simulated dataset and two real datasets, and applied the Coherent Point Drift (CPD) algorithm [32] to align embeddings from the five foundational models, with and without DRIFT. We also evaluated the results using two specialized methods, PASTE and PASTE2. We formulated all the tasks as zero-shot problems, with no re-training or fine-tuning. After alignment, we measure the similarity between the transcriptomics profiles of the paired cells using the metrics described in the Methods section.

#### Simulations

To assess DRIFT’s robustness to noise, we simulated ten paired ST slices from a manually aligned 10xHSI (Visium human ovarian cancer) pair, varying the pseudocount level (*δ* from 0.5 to 5) to introduce increased expression noise (as done in [27, 20]). We compared DRIFT and baseline alignments using translation error (lower is better), Pearson Correlation, and Kendall’s *τ* (higher is better).

Across all settings, DRIFT consistently reduces translation error for all scRNA-seq foundation models. Geneformer, scFoundation, and scGPT show the most substantial gains, with typical reductions ranging from 30% to 77% across most *δ* values. ST foundation models also benefit from DRIFT, where Nicheformer and Loki show reliable improvements for very noisy samples, with reductions of at least 40%.

This positive improvement is further supported by systematically higher PCC values (0.5% to 6% gains) and higher Kendall’s *τ* (0.2% to 4%). PASTE and PASTE2, on the other hand, have higher translation error and lower PCC and Kendall’s *τ* (Figure 3 and Supplementary Figure S23).

**Fig. 3.**
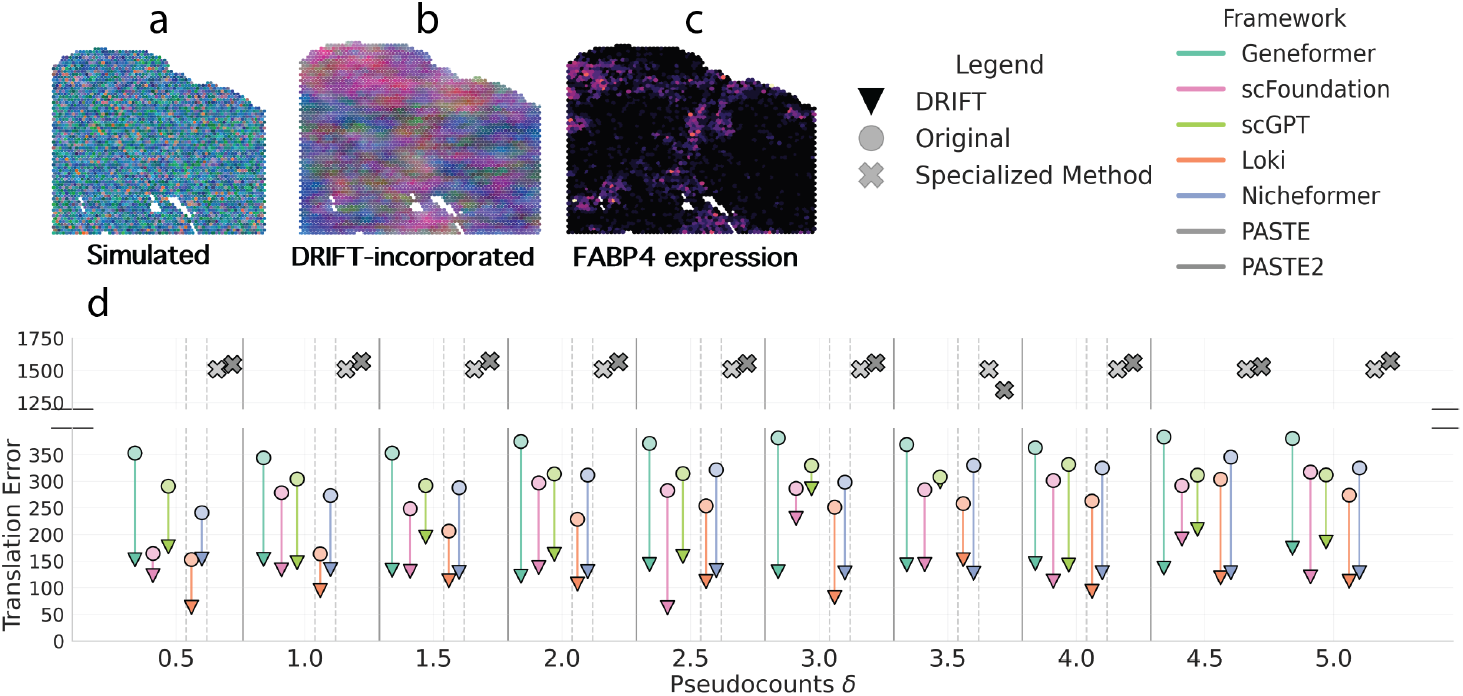
Simulated Data Alignment Results. a–c. PCA visualizations of the original and DRIFT-incorporated embeddings for simulated data at pseudocount 4.0, along with the corresponding FABP4 expression patterns. DRIFT produces embeddings with clearer and more coherent spatial organization, recovering biologically meaningful structure associated with ovarian cancer development. d. Translation error across simulated datasets at different values of pseudocounts, demonstrating improved alignment fidelity after applying DRIFT.

Next, since Geneformer showed the greatest percentage reduction in translation error, we performed principal component analysis (PCA) on the original and DRIFT-incorporated Geneformer embeddings at a pseudocount of 4.0 for illustration. While the original embeddings appeared noisy, DRIFT embeddings show improved spatial organization consistent with important marker gene patterns in ovarian cancer (see Figure 3a-c). Altogether, these results confirm that DRIFT not only reduces spatial reconstruction error but also recovers the underlying spatial gene expression patterns.

#### Mouse embryonic dataset

We then applied DRIFT to reconstruct three-dimensional tissue organization using the StereoME (STEREO-seq) mouse embryo dataset, which provides spatial maps across early organogenesis. We aligned four consecutive E9.5 sections (Slice 1→2, 2→3, and 3→4) and reconstructed the resulting embeddings in 3D space. Because ground truth is unavailable in this setting, we evaluated reconstruction quality through visualization and the alignment score. Visually, only the original scFoundation embeddings produce reasonably coherent alignments, whereas the other foundation models frequently yield inconsistent mappings, such as inversions or rotations, highlighting the difficulty of this alignment task (Figure 4, Supplementary Figure S24, and Supplementary Section S10). Incorporating DRIFT noticeably improves performance across all models in most scenarios, making 3D reconstructions more stable and biologically consistent.

**Fig. 4.**
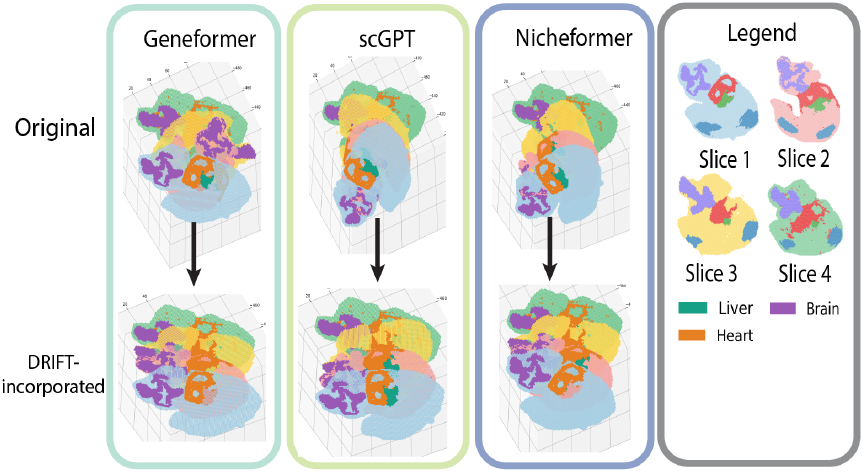
Select Mouse Embryo Alignment Results. **First row:** alignments obtained using the original embeddings of Geneformer, scFoundation, and Nichefomer. **Second row:** alignments obtained using respective DRIFT-incorporated embeddings. Across slices, DRIFT yields stable and spatially coherent embryonic alignments, resulting in higher-fidelity three-dimensional tissue reconstructions. Full results are available in Supplementary Figure S24.

Quantitatively, the scRNA-seq foundation models exhibit strong gains under DRIFT, with Geneformer and scGPT showing large average improvements (up to 46.5% to 73.6%, respectively), reflecting DRIFT’s ability to correct poor baseline alignments (Supplementary Section S10). scFoundation, which already produces relatively strong alignments, also shows improved performance (averaging ≈ 10% increase), indicating that DRIFT can correct high-quality embeddings. The ST foundation models, especially Nicheformer, also benefit, particularly in Sllice 3→4, with an average gain of 17.8% while Loki’s performance increases by over 24.1% on average. As shown in Supplementary Section S10, some embeddings achieve higher alignment scores than PASTE across slices and remain competitive with, or even outperform, PASTE2 only when combined with DRIFT, which is required for these models to achieve their strongest alignment performance.

#### Human small intestine dataset

Finally, to further validate DRIFT’s alignment capability in real-world contexts, we aligned three sets of four adjacent human small intestine sections manually annotated by experts onto (10xHSI) from the same specimen. Across all experiments, DRIFT improves alignments for every foundation model evaluated. The magnitude of the gain differs across frameworks: scFoundation shows the strongest and most stable increase, with several experiments exhibiting significant boosts (up to 33%). Geneformer and scGPT also demonstrate reliable improvements (7% to 17%), suggesting that DRIFT effectively enhances context-aware transcriptomic embeddings (Figure 5). ST foundation models show similar trends, with Nicheformer benefiting from substantial improvements up to 22% while Loki exhibits smaller but still positive gains across most experiments (between 1 and 5%). DRIFT-incorporated foundation models perform competitively in eight of the nine experiments – outperforming PASTE and PASTE2 in six cases and matching their performance in an additional two, indicating that this generalizable diffusion refinement can outperform specialized alignment tools even under challenging structural variation.

**Fig. 5.**
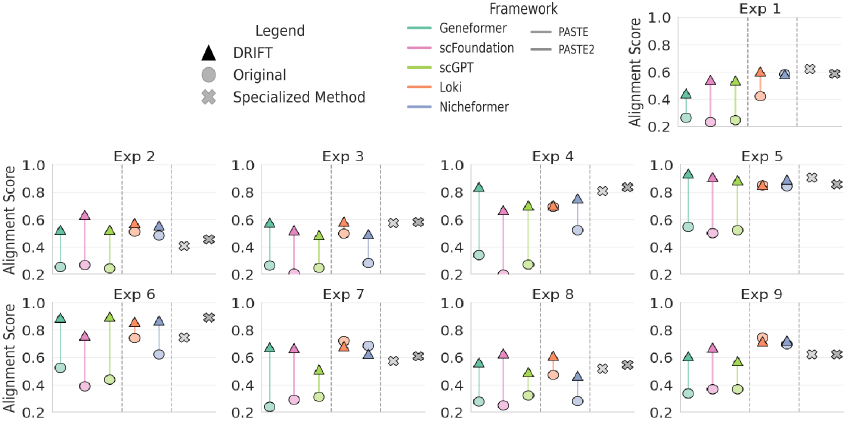
Human Small Intestine Alignment Results. Foundation models substantially improve alignment quality across human small intestine sections and become competitive with PASTE/PASTE2 when paired with DRIFT.

Altogether, DRIFT denoises spatial embeddings and reconstructs meaningful biological structure, yielding more accurate and coherent alignments.

### DRIFT enhances clustering performance

Finally, we evaluated DRIFT’s performance on the unsupervised clustering task. We formulated the task as a zero-shot problem, with no re-training or fine-tuning. We used the pretrained weights of the foundation models (with and without DRIFT inputs) to obtain cell/spot embeddings and subsequently clustered them using the mclust algorithm [35]. We evaluated all methods on the five datasets from the annotation task. The five datasets were 10xHPC (10x human prefrontal cortex), MERHH (MERFISH human heart), MERMPH (MERFISH mouse preoptic hypothalamus), MERMB (MERFISH mouse brain), and MERMBA (MERFISH mouse brain). The ground truth for clustering is cell-type labels annotated by experts.

We clustered each slice independently and quantified the performance of the five foundation models and one specialized method (GraphST) with ARI as the evaluation metric.

As displayed in Table 2, DRIFT improves the zero-shot performance of all scRNA-seq foundation models. Quantitatively, the mean ARI scores rose substantially from 0.0778 to 0.2681 for Geneformer, 0.2558 to 0.5233 for scFoundation, and 0.2983 to 0.4470 for scGPT. These results further demonstrate that the diffused inputs generated by DRIFT better capture biologically meaningful structure, enabling foundation models to access richer spatial–molecular information in a zero-shot setting.

**Table 2.**
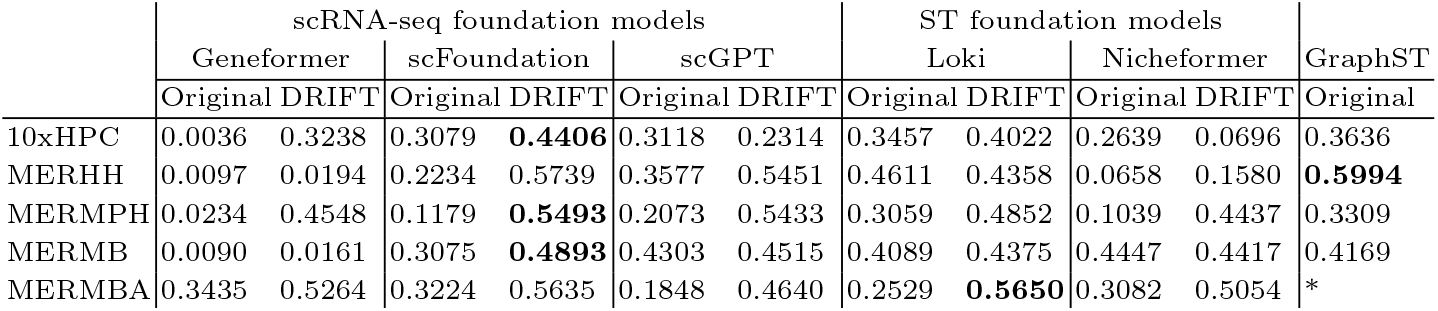
Clustering Results. The table lists the mean zero-shot ARI performance of every foundation model and its DRIFT-incorporated versions on each dataset. These results show that the best-performing foundation models overall are DRIFT-incorporated scGPT, DRIFT-incorporated scFoundation, and DRIFT-incorporated Loki. *-GraphST did not run on the MERMBA dataset.

We also see improved performance for ST foundation models. The mean ARI scores rose from 0.3549 to 0.4651 for Loki, and from 0.2372 to 0.3236 for Nicheformer, confirming the effectiveness of DRIFT even on models trained with spatial data.

Interestingly, we observe that scFoundation, scGPT, and Loki models with DRIFT input outperformed GraphST’s mean ARI score of 0.4277, achieving mean ARIs of 0.5132, 0.4428, and 0.4401, respectively, on the four datasets for which GraphST could be successfully executed. Note that GraphST requires re-training on new samples, whereas foundation models with DRIFT input are run in a zero-shot setting. This result demonstrates that DRIFT effectively bridges the performance gap between general-purpose foundation models and specialized ST methods, allowing previously underperforming foundation models to achieve or exceed ST-specific benchmarks. We also observe that the performance of some of the foundation models can be further improved with few-shot training, as outlined in Supplementary Section S11.

## Conclusion

We introduce DRIFT, a graph diffusion–based framework that integrates spatial context into the inputs of pretrained foundation models. DRIFT-incorporated foundation models improve performance across three ST tasks — annotation, alignment, and clustering — without requiring architectural changes or retraining.

DRIFT is a simple, effective, and accessible method for coupling pretrained scRNA-seq foundation models with ST data and performing diffusion-based denoising for ST foundational models. By combining pretrained embeddings with diffusion-enhanced information, DRIFT leverages existing models and extends them to a unified framework for spatially aware representations of tissues and developmental organization.

