## Supplementary File for "Diffusion-based Representation Integration for Foundation Models Improves Spatial Transcriptomics Analysis"

### Supplementary Information

#### S1 Graph Construction Comparison

To identify the optimal graph topology for DRIFT, we evaluated two construction techniques:

- **K-Nearest Neighbors** - For this graph construction technique, we connected each node to the k spatially closest nodes based on Euclidean distance. Then we converted all edges to undirected, ensuring that every node has a minimum degree of $k$ (though some may have a higher degree). We refer to this construction as Kk, where k denotes the number of closest neighbors (e.g., K3 = 3-nearest neighbors).
- **Threshold-based Method** - For this technique, we determined the spatial Euclidean distance threshold that yields an *average* of n neighbors across the graph. Unlike the neighbor-based approach, this method allows the number of connections per node to vary significantly based on local tissue density. We refer to this construction as Tn, where n denotes the number of average neighbors (e.g., T3 = Threshold-based construction with an average of 3 neighbors).

As observed in Figures S1 and S2, the K3 construction consistently yielded superior performance on the annotation task for scGPT (the best-performing annotation foundation model), demonstrating robustness across different ST platform resolutions. Therefore, we use K3 as our graph construction technique of choice.

#### S2 Diffusion and Denoising Methods Comparison

We compared our heat kernel diffusion method to multiple graph diffusion techniques on the cell annotation task. The graph diffusion techniques are:

- **Random Walk Diffusion** - Let $A$ be the adjacency matrix of the graph. The we calculate the degree matrix $D$ as:

  $$D_{ii} = \sum_j A_{ij} \tag{1}$$

  We then calculate the transition matrix $P$ for a random walk as:

  $$P = D^{-1}A \tag{2}$$

  Finally, given a the input gene expression matrix $x_0$, we calculate the diffused output matrix $x_k$ as:

  $$x_k = P^k x_o \tag{3}$$

  Where k is a positive integer hyperparameter that determines the diffusion strength of the method. Higher $k$ values result in smoothing across larger spatial distances. Since k can only take positive integer values, it is limited in how much it can be tuned relative to the heat kernel.
- **Mean** - For every node in mean diffusion, we take the gene expression vectors of the node and all its neighbors and return the mean expression as the diffused output.
- **Median** - For every node in median diffusion, we take the gene expression vectors of the node and all its neighbors and return the median expression as the diffused output.
- **Spectral Gaussian Filter** - The spectral Gaussian filter is a type of low-pass filter calculated through the eigenvalue decomposition. Let $L$ be the normalized Laplacian of the graph.

  $$L = U\Lambda U^{-1} = U\Lambda U^{\top} \tag{4}$$

  We then apply the Gaussian filter $g(l)$ on the eigenvalues $l$. The filter is defined as:

  $$g(l) = e^{\frac{-l^2}{2\sigma^2}} \tag{5}$$

  where $\sigma$ is the standard deviation of the Gaussian, which we set to 1.

Table S1: Graph shuffling results in reduced cell-annotation model f1-score performance, indicating local spatial context is necessary for DRIFT. Shuffled Diffusion represents shuffling the neighbors in the graph and then performing diffusion. Higher values correspond to better performance.

| | | 10xHPC | MERHH | MERMPH | MERMB | MERMBA |
|---|---|---|---|---|---|---|
| Geneformer | DRIFT | 0.6085 | 0.671 | 0.6858 | 0.4286 | 0.4745 |
| | Shuffled Diffusion | 0.1315 | 0.059 | 0.0872 | 0.0718 | 0.0627 |
| scFoundation | DRIFT | 0.6017 | 0.6357 | 0.7178 | 0.4899 | 0.5829 |
| | Shuffled Diffusion | 0.1315 | 0.0872 | 0.059 | 0.0718 | 0.0627 |
| scGPT | DRIFT | 0.661 | 0.756 | 0.765 | 0.572 | 0.703 |
| | Shuffled Diffusion | 0.1133 | 0.045 | 0.06 | 0.066 | 0.042 |
| Loki | DRIFT | 0.6558 | 0.6731 | 0.7094 | 0.3794 | 0.6141 |
| | Shuffled Diffusion | 0.1351 | 0.0562 | 0.0702 | 0.0777 | 0.0421 |
| Nicheformer | DRIFT | 0.5659 | 0.6523 | 0.6754 | 0.4579 | 0.6068 |
| | Shuffled Diffusion | 0.1238 | 0.0329 | 0.0717 | 0.0707 | 0.0461 |

Table S2: Neighborhood composition prediction shows closer prediction to ground truth distributions with DRIFT. Original refers to performing the neighborhood task without any diffusion. Lower values correspond to better performance.

| | | MERHH | MERHH | MERMPH | MERMB | MERMBA |
|---|---|---|---|---|---|---|
| Geneformer | DRIFT | 0.0501 | 0.0567 | 0.0577 | 0.0308 | 0.0353 |
| | Original | 0.0734 | 0.0565 | 0.1029 | 0.0592 | 0.0675 |
| scFoundation | DRIFT | 0.023154 | 0.016641 | 0.033013 | 0.018502 | 0.026804 |
| | Original | 0.043738 | 0.027556 | 0.054934 | 0.031691 | 0.054698 |
| scGPT | DRIFT | 0.0634 | 0.0196 | 0.0403 | 0.0333 | 0.0402 |
| | Original | 0.0526 | 0.0302 | 0.0606 | 0.0413 | 0.0700 |
| Loki | DRIFT | 0.029762 | 0.013176 | 0.036412 | 0.027431 | 0.030464 |
| | Original | 0.049638 | 0.027323 | 0.056532 | 0.037651 | 0.056513 |
| Nicheformer | DRIFT | 0.038249 | 0.019111 | 0.036169 | 0.019549 | 0.032661 |
| | Original | 0.053462 | 0.034545 | 0.058315 | 0.0292 | 0.057233 |

- **Spectral Low-Pass Filter** - The spectral Low-Pass filter is a hard low-pass filter where any eigenvalues above a threshold are set to 0. Since we calculate the eigenvalues for a normalized Laplacian, the threshold is set at 0.5.
- **GraphST** - GraphST denoises data by aggregating spatial graph-based features, reconstructing gene expression, and utilizing contrastive learning to reinforce consistent spatial signals. [1], [2]

As seen in Figures S3-S16, the heat kernel shows an overall higher performance for top-performing foundation models, thus setting it as our choice of graph diffusion.

### S3 Spatial Awareness Experiments

To evaluate the extent to which DRIFT-incorporated embeddings capture local spatial information, we conducted two complementary experiments. First, we extended the cell type annotation task by perturbing local spatial structure through randomized graph connectivity. Second, we assessed neighborhood awareness by formulating a neighboring-cell-type composition task, in which models predict the proportions of cell types within predefined spatial radii around each cell, similar to Schaar *et al.*[3]

In the first experiment, we assessed sensitivity to local spatial structure by randomizing graph connectivity prior to cell type annotation. As observed in Table S1, across datasets and foundation models, this perturbation led to a noticeable decline in annotation performance, indicating that accurate predictions depend on preserved spatial neighborhood information.

[1] Yahui Long, Kok Siong Ang, Mengwei Li, Kian Long Kelvin Chong, Raman Sethi, Chengwei Zhong, Hang Xu, Zhiwei Ong, Karishma Sachaphibulkij, Ao Chen, et al. Spatially informed clustering, integration, and deconvolution of spatial transcriptomics with graphst. *Nature Communications*, 14(1):1155, 2023

[2] Haiyue Wang, Peng Gao, Shaoqing Feng, and Xiaoke Ma. Denoising spatially resolved transcriptomics with consistency of heterogeneous spatial coordinates, transcription, and morphology. *Briefings in Bioinformatics*, 26(5):bbaf528, 2025

[3] Anna C Schaar, Alejandro Tejada-Lapuerta, Giovanni Palla, Robert Gutgesell, Lennard Halle, Mariia Minaeva, Larsen Vornholz, Leander Dony, Francesca Drummer, Mojtaba Bahrami, et al. Nicheformer: a foundation model for single-cell and spatial omics. *bioRxiv*, pages 2024–04, 2024

In the second experiment, we evaluated neighborhood awareness by predicting the composition of cell types within predefined spatial radii around each cell, with performance measured using KL divergence between predicted and true neighborhood distributions. Specifically, given a true neighborhood composition $P = (p_1, \ldots, p_C)$ and a predicted composition $Q = (q_1, \ldots, q_C)$ over $C$ cell types, we compute $\mathrm{KL}(P \| Q) = \sum_{c=1}^{C} p_c \log \frac{p_c}{q_c}$, where lower values indicate closer agreement between predicted and true neighborhood compositions. Across datasets and foundation models, DRIFT-incorporated embeddings consistently achieved lower KL divergence compared to baseline embeddings, indicating more accurate recovery of local cell type composition (Table S2).

Together, these two experiments demonstrate that DRIFT explicitly encodes local spatial context, with performance degrading under disrupted spatial structure but yielding more accurate neighborhood-level representations when spatial relationships are preserved.

### S4 Geneformer Performance

We compared all three Geneformer models on the cell annotation task. As observed in Figures S11-S16, Geneformer V2 with 316 million parameters performed the best. Therefore, we use it as our choice of Geneformer.

### S5 Foundation Model Technical Explanation

To ensure consistency with downstream foundation models trained on human gene identifiers, mouse datasets were mapped to their corresponding human orthologs using Ensembl BioMart. For each mouse gene, the associated human ortholog symbol and Ensembl gene identifier were retrieved. Genes without valid ortholog mappings were discarded, and duplicate mappings were resolved by retaining a single representative per mouse gene. The dataset was subsequently restricted to genes with valid orthologs, with human gene symbols used as primary identifiers.

The processed datasets were subsequently used as inputs to the DRIFT framework prior to integration with foundation models. Because each foundation model operates on a predefined gene vocabulary, genes not represented in a model's native vocabulary were handled according to model-specific preprocessing strategies to ensure compatible input dimensionality. Different normalization and selection procedures were applied: Loki retained the top 50 most highly expressed genes per cell, scFoundation accommodated missing genes by assigning zero values, Nicheformer and Geneformer ranked genes within each cell based on expression magnitude, and scGPT discretized gene expression values into categorical bins. These procedures ensured consistency between the input data and each model's representation while preserving relative expression levels.

### S6 Hyperparameter Tuning $t$

To test which value of $t$ performed best, we ran all methods across all tasks with the following values of $t$ - $10^{-5}$, $10^{-4}$, $10^{-3}$, $10^{-2}$, $10^{-1}$, 1, 2, 5, 10, and 100. The performance of different values of the hyperparameter $t$ in each task is illustrated in the following images:

- **Cell Annotation** - Figures S3-S12
- **Slice Alignment** - Tables S5 to S9
- **Clustering** - Figures S17-S21

Across all tasks, we note that $t \in \{1, 2, 5, 10\}$ tend to yield the best results.

For annotation and clustering, $t = 5$ performs well overall and is therefore chosen as the default value for these tasks.

For slice alignment, we select the hyperparameter that yields the highest Pearson correlation. Because this metric does not rely on labels, the tuning procedure can be applied in any setting.

### S7 Hyperparameter Tuning Probes for Few-Shot Learning

To hyperparameter-tune the three-layered multilayer perceptron (MLP) foundation model probes that we train for few-shot learning, we employed Optuna for 10 trials. The range of values that we searched over is shown in Table S3.

Table S3: The range of values used for each hyperparameter during hyperparameter tuning.

| **Hyperparameter** | **Values** |
|---|---|
| Hidden Size of Layer 1 | 10 - input embedding size |
| Hidden Size of Layer 2 | 10 - 256 |
| Learning rate | $10^{-5}$ - $10^{-1}$ |
| Weight Decay | $10^{-6}$ - $10^{-3}$ |
| Dropout | 0.1 - 0.6 |
| Use Class Weights for Loss | True/False |

### S8 Hyperparameter Tuning Specialized Methods

#### 5.1 STELLAR

For STELLAR, we also used Optuna for 10 trials to hyperparameter-tune the method. The range of values that we searched over is shown in Table S4.

Table S4: The range of values used for each hyperparameter during hyperparameter tuning.

| **Hyperparameter** | **Values** |
|---|---|
| Learning rate | $10^{-5}$ - $10^{-1}$ |
| Weight Decay | $5 * 10^{-3}$ - $5 * 10^{-1}$ |
| Seed | 1 - 20 |

#### 5.2 PASTE and PASTE2

We tuned the hyperparameter $\alpha \in \{0.1 - 0.9\}$ with a step size of 0.1 for both PASTE and PASTE2, selecting the configuration that achieved the highest Pearson correlation.

#### 5.3 GraphST

As mentioned in Section 3.3 of the main text, since clustering is an unsupervised task, we did not perform any hyperparameter tuning for the foundation models and GraphST.

### S9 Annotation Accuracy Results

Figure S22 demonstrates DRIFT's improved annotation accuracy performance across foundation models and datasets. These performances are also compared to a state-of-the-art annotation method, STELLAR.

### S10 Alignment Results

Tables S5 to S9 show alignment results on mouse embryonic reconstruction for all foundation models with and without DRIFT.

Table S5: Geneformer's performance on mouse embryonic reconstruction

| Experiment | Foundation model | Method | LTARI | Pearson | Kendall |
|---|---|---|---|---|---|
| E95_S1 → E95_S2 | Geneformer | Original | 0.0585 | 0.8243 | 0.3246 |
| E95_S1 → E95_S2 | Geneformer | Diffusion_5.0 | 0.1258 | 0.8532 | 0.3456 |
| E95_S1 → E95_S2 | Geneformer | Diffusion_2.0 | 0.0855 | 0.8478 | 0.3408 |
| E95_S1 → E95_S2 | Geneformer | Diffusion_100.0 | 0.1437 | 0.8501 | 0.3433 |
| E95_S1 → E95_S2 | Geneformer | Diffusion_10.0 | 0.1292 | 0.851 | 0.3441 |
| E95_S1 → E95_S2 | Geneformer | Diffusion_1.0 | 0.0837 | 0.8468 | 0.3402 |
| E95_S1 → E95_S2 | Geneformer | Diffusion_0.1 | 0.1436 | 0.8614 | 0.3497 |
| E95_S1 → E95_S2 | Geneformer | Diffusion_0.01 | 0.1425 | 0.8607 | 0.3491 |
| E95_S1 → E95_S2 | Geneformer | Diffusion_0.001 | 0.0577 | 0.8273 | 0.3251 |
| E95_S1 → E95_S2 | Geneformer | Diffusion_0.0001 | 0.0432 | 0.8301 | 0.3253 |
| E95_S1 → E95_S2 | Geneformer | Diffusion_0.00001 | 0.0518 | 0.831 | 0.3248 |
| E95_S2 → E95_S3 | Geneformer | Original | 0.0858 | 0.8185 | 0.3239 |
| E95_S2 → E95_S3 | Geneformer | Diffusion_5.0 | 0.1562 | 0.8462 | 0.3366 |
| E95_S2 → E95_S3 | Geneformer | Diffusion_2.0 | 0.1622 | 0.8462 | 0.3396 |
| E95_S2 → E95_S3 | Geneformer | Diffusion_100.0 | 0.1924 | 0.8447 | 0.3402 |
| E95_S2 → E95_S3 | Geneformer | Diffusion_10.0 | 0.2017 | 0.8477 | 0.3402 |
| E95_S2 → E95_S3 | Geneformer | Diffusion_1.0 | 0.1718 | 0.8501 | 0.3423 |
| E95_S2 → E95_S3 | Geneformer | Diffusion_0.1 | 0.1948 | 0.8517 | 0.3436 |
| E95_S2 → E95_S3 | Geneformer | Diffusion_0.01 | 0.1855 | 0.8504 | 0.3431 |
| E95_S2 → E95_S3 | Geneformer | Diffusion_0.001 | 0.2151 | 0.8561 | 0.3413 |
| E95_S2 → E95_S3 | Geneformer | Diffusion_0.0001 | 0.1734 | 0.8525 | 0.3373 |
| E95_S2 → E95_S3 | Geneformer | Diffusion_0.00001 | 0.095 | 0.8113 | 0.3232 |
| E95_S3 → E95_S4 | Geneformer | Original | 0.1217 | 0.8138 | 0.3256 |
| E95_S3 → E95_S4 | Geneformer | Diffusion_5.0 | 0.1603 | 0.849 | 0.3409 |
| E95_S3 → E95_S4 | Geneformer | Diffusion_2.0 | 0.1068 | 0.7835 | 0.329 |
| E95_S3 → E95_S4 | Geneformer | Diffusion_100.0 | 0.1129 | 0.8217 | 0.3348 |
| E95_S3 → E95_S4 | Geneformer | Diffusion_10.0 | 0.1628 | 0.8497 | 0.3416 |
| E95_S3 → E95_S4 | Geneformer | Diffusion_1.0 | 0.1225 | 0.8176 | 0.3341 |
| E95_S3 → E95_S4 | Geneformer | Diffusion_0.1 | 0.2557 | 0.8728 | 0.3546 |
| E95_S3 → E95_S4 | Geneformer | Diffusion_0.01 | 0.2034 | 0.8619 | 0.3513 |
| E95_S3 → E95_S4 | Geneformer | Diffusion_0.001 | 0.1369 | 0.8049 | 0.3321 |
| E95_S3 → E95_S4 | Geneformer | Diffusion_0.0001 | 0.1139 | 0.8129 | 0.3252 |
| E95_S3 → E95_S4 | Geneformer | Diffusion_0.00001 | 0.1245 | 0.8167 | 0.3264 |

Table S6: scFoundation's performance on mouse embryonic reconstruction

| **Experiment** | **Foundation model** | **Method** | **LTARI** | **Pearson** | **Kendall** |
|---|---|---|---|---|---|
| E95_S1 → E95_S2 | scfoundation | Original | 0.1571 | 0.869 | 0.3526 |
| E95_S1 → E95_S2 | scfoundation | Diffusion_5.0 | 0.2116 | 0.8762 | 0.3567 |
| E95_S1 → E95_S2 | scfoundation | Diffusion_2.0 | 0.0592 | 0.8408 | 0.3247 |
| E95_S1 → E95_S2 | scfoundation | Diffusion_100.0 | 0.162 | 0.8602 | 0.3503 |
| E95_S1 → E95_S2 | scfoundation | Diffusion_10.0 | 0.2283 | 0.8753 | 0.3586 |
| E95_S1 → E95_S2 | scfoundation | Diffusion_1.0 | 0.1604 | 0.864 | 0.3528 |
| E95_S1 → E95_S2 | scfoundation | Diffusion_0.1 | 0.246 | 0.8796 | 0.3603 |
| E95_S1 → E95_S2 | scfoundation | Diffusion_0.01 | 0.2093 | 0.8724 | 0.3567 |
| E95_S1 → E95_S2 | scfoundation | Diffusion_0.001 | 0.2389 | 0.8801 | 0.3595 |
| E95_S1 → E95_S2 | scfoundation | Diffusion_0.0001 | 0.2374 | 0.8804 | 0.3597 |
| E95_S1 → E95_S2 | scfoundation | Diffusion_0.00001 | 0.2383 | 0.8804 | 0.3599 |
| E95_S2 → E95_S3 | scfoundation | Original | 0.2074 | 0.8523 | 0.3438 |
| E95_S2 → E95_S3 | scfoundation | Diffusion_5.0 | 0.1946 | 0.8467 | 0.3415 |
| E95_S2 → E95_S3 | scfoundation | Diffusion_2.0 | 0.1894 | 0.8492 | 0.3416 |
| E95_S2 → E95_S3 | scfoundation | Diffusion_100.0 | 0.1723 | 0.8429 | 0.3357 |
| E95_S2 → E95_S3 | scfoundation | Diffusion_10.0 | 0.2061 | 0.8492 | 0.339 |
| E95_S2 → E95_S3 | scfoundation | Diffusion_1.0 | 0.1866 | 0.8477 | 0.3416 |
| E95_S2 → E95_S3 | scfoundation | Diffusion_0.1 | 0.1927 | 0.8458 | 0.3412 |
| E95_S2 → E95_S3 | scfoundation | Diffusion_0.01 | 0.1888 | 0.8473 | 0.3407 |
| E95_S2 → E95_S3 | scfoundation | Diffusion_0.001 | 0.1804 | 0.8475 | 0.3421 |
| E95_S2 → E95_S3 | scfoundation | Diffusion_0.0001 | 0.1793 | 0.8467 | 0.3422 |
| E95_S2 → E95_S3 | scfoundation | Diffusion_0.00001 | 0.1797 | 0.8469 | 0.3423 |
| E95_S3 → E95_S4 | scfoundation | Original | 0.1732 | 0.8496 | 0.3426 |
| E95_S3 → E95_S4 | scfoundation | Diffusion_5.0 | 0.1538 | 0.8472 | 0.3452 |
| E95_S3 → E95_S4 | scfoundation | Diffusion_2.0 | 0.1951 | 0.8593 | 0.3505 |
| E95_S3 → E95_S4 | scfoundation | Diffusion_100.0 | 0.1662 | 0.849 | 0.3432 |
| E95_S3 → E95_S4 | scfoundation | Diffusion_10.0 | 0.1319 | 0.8258 | 0.338 |
| E95_S3 → E95_S4 | scfoundation | Diffusion_1.0 | 0.2154 | 0.864 | 0.3522 |
| E95_S3 → E95_S4 | scfoundation | Diffusion_0.1 | 0.1341 | 0.8153 | 0.3276 |
| E95_S3 → E95_S4 | scfoundation | Diffusion_0.01 | 0.1386 | 0.8283 | 0.3379 |
| E95_S3 → E95_S4 | scfoundation | Diffusion_0.001 | 0.1359 | 0.8498 | 0.3422 |
| E95_S3 → E95_S4 | scfoundation | Diffusion_0.0001 | 0.1325 | 0.849 | 0.3417 |
| E95_S3 → E95_S4 | scfoundation | Diffusion_0.00001 | 0.1326 | 0.8492 | 0.3417 |

Table S7: scGPT's performance on mouse embryonic reconstruction

| Experiment | Foundation model | Method | LTARI | Pearson | Kendall |
|---|---|---|---|---|---|
| E95_S1 → E95_S2 | scgpt | Original | 0.0926 | 0.8493 | 0.3347 |
| E95_S1 → E95_S2 | scgpt | Diffusion_5.0 | 0.223 | 0.8783 | 0.3592 |
| E95_S1 → E95_S2 | scgpt | Diffusion_2.0 | 0.2316 | 0.8775 | 0.3601 |
| E95_S1 → E95_S2 | scgpt | Diffusion_100.0 | 0.254 | 0.8799 | 0.3604 |
| E95_S1 → E95_S2 | scgpt | Diffusion_10.0 | 0.23 | 0.8781 | 0.3596 |
| E95_S1 → E95_S2 | scgpt | Diffusion_1.0 | 0.214 | 0.8776 | 0.3581 |
| E95_S1 → E95_S2 | scgpt | Diffusion_0.1 | 0.2144 | 0.8767 | 0.3569 |
| E95_S1 → E95_S2 | scgpt | Diffusion_0.01 | 0.218 | 0.8771 | 0.3585 |
| E95_S1 → E95_S2 | scgpt | Diffusion_0.001 | 0.2126 | 0.8783 | 0.3581 |
| E95_S1 → E95_S2 | scgpt | Diffusion_0.0001 | 0.2478 | 0.8813 | 0.3603 |
| E95_S1 → E95_S2 | scgpt | Diffusion_0.00001 | 0.1997 | 0.8679 | 0.3545 |
| E95_S2 → E95_S3 | scgpt | Original | 0.1668 | 0.8471 | 0.339 |
| E95_S2 → E95_S3 | scgpt | Diffusion_5.0 | 0.195 | 0.8518 | 0.3435 |
| E95_S2 → E95_S3 | scgpt | Diffusion_2.0 | 0.2013 | 0.8517 | 0.3437 |
| E95_S2 → E95_S3 | scgpt | Diffusion_100.0 | 0.1958 | 0.8476 | 0.3415 |
| E95_S2 → E95_S3 | scgpt | Diffusion_10.0 | 0.1925 | 0.8493 | 0.3428 |
| E95_S2 → E95_S3 | scgpt | Diffusion_1.0 | 0.2 | 0.8523 | 0.3447 |
| E95_S2 → E95_S3 | scgpt | Diffusion_0.1 | 0.1961 | 0.8526 | 0.3441 |
| E95_S2 → E95_S3 | scgpt | Diffusion_0.01 | 0.1929 | 0.8504 | 0.3428 |
| E95_S2 → E95_S3 | scgpt | Diffusion_0.001 | 0.1842 | 0.85 | 0.3431 |
| E95_S2 → E95_S3 | scgpt | Diffusion_0.0001 | 0.1907 | 0.8518 | 0.3448 |
| E95_S2 → E95_S3 | scgpt | Diffusion_0.00001 | 0.1892 | 0.8506 | 0.3427 |
| E95_S3 → E95_S4 | scgpt | Original | 0.08 | 0.7872 | 0.3255 |
| E95_S3 → E95_S4 | scgpt | Diffusion_5.0 | 0.1271 | 0.8367 | 0.3393 |
| E95_S3 → E95_S4 | scgpt | Diffusion_2.0 | 0.2429 | 0.8716 | 0.3539 |
| E95_S3 → E95_S4 | scgpt | Diffusion_100.0 | 0.1382 | 0.7928 | 0.3288 |
| E95_S3 → E95_S4 | scgpt | Diffusion_10.0 | 0.1718 | 0.8538 | 0.3481 |
| E95_S3 → E95_S4 | scgpt | Diffusion_1.0 | 0.2548 | 0.8783 | 0.3545 |
| E95_S3 → E95_S4 | scgpt | Diffusion_0.1 | 0.1418 | 0.8486 | 0.3436 |
| E95_S3 → E95_S4 | scgpt | Diffusion_0.01 | 0.1077 | 0.8289 | 0.3261 |
| E95_S3 → E95_S4 | scgpt | Diffusion_0.001 | 0.1228 | 0.8317 | 0.3256 |
| E95_S3 → E95_S4 | scgpt | Diffusion_0.0001 | 0.1511 | 0.8552 | 0.3422 |
| E95_S3 → E95_S4 | scgpt | Diffusion_0.00001 | 0.1325 | 0.8342 | 0.3394 |

Table S8: Loki's performance on mouse embryonic reconstruction

| Experiment | Foundation model | Method | LTARI | Pearson | Kendall |
|---|---|---|---|---|---|
| E95_S1 → E95_S2 | Loki | Original | 0.0683 | 0.832 | 0.3282 |
| E95_S1 → E95_S2 | Loki | Diffusion_5.0 | 0.2153 | 0.8722 | 0.3576 |
| E95_S1 → E95_S2 | Loki | Diffusion_2.0 | 0.1618 | 0.864 | 0.3526 |
| E95_S1 → E95_S2 | Loki | Diffusion_100.0 | 0.163 | 0.867 | 0.3534 |
| E95_S1 → E95_S2 | Loki | Diffusion_10.0 | 0.2236 | 0.8754 | 0.3581 |
| E95_S1 → E95_S2 | Loki | Diffusion_1.0 | 0.1844 | 0.8696 | 0.3546 |
| E95_S1 → E95_S2 | Loki | Diffusion_0.1 | 0.0773 | 0.8326 | 0.319 |
| E95_S1 → E95_S2 | Loki | Diffusion_0.01 | 0.0852 | 0.8424 | 0.3296 |
| E95_S1 → E95_S2 | Loki | Diffusion_0.001 | 0.0856 | 0.8411 | 0.3305 |
| E95_S1 → E95_S2 | Loki | Diffusion_0.0001 | 0.0854 | 0.8412 | 0.3305 |
| E95_S1 → E95_S2 | Loki | Diffusion_0.00001 | 0.0855 | 0.8413 | 0.3305 |
| E95_S2 → E95_S3 | Loki | Original | 0.1638 | 0.843 | 0.3365 |
| E95_S2 → E95_S3 | Loki | Diffusion_5.0 | 0.1729 | 0.8438 | 0.3387 |
| E95_S2 → E95_S3 | Loki | Diffusion_2.0 | 0.0846 | 0.8434 | 0.3229 |
| E95_S2 → E95_S3 | Loki | Diffusion_100.0 | 0.169 | 0.8494 | 0.3334 |
| E95_S2 → E95_S3 | Loki | Diffusion_10.0 | 0.2155 | 0.8548 | 0.3378 |
| E95_S2 → E95_S3 | Loki | Diffusion_1.0 | 0.1853 | 0.8433 | 0.337 |
| E95_S3 → E95_S4 | Loki | Original | 0.2777 | 0.8826 | 0.359 |
| E95_S3 → E95_S4 | Loki | Diffusion_5.0 | 0.2718 | 0.8795 | 0.3564 |
| E95_S3 → E95_S4 | Loki | Diffusion_2.0 | 0.1921 | 0.8612 | 0.3509 |
| E95_S3 → E95_S4 | Loki | Diffusion_100.0 | 0.184 | 0.8465 | 0.3464 |
| E95_S3 → E95_S4 | Loki | Diffusion_10.0 | 0.2547 | 0.8765 | 0.3549 |
| E95_S3 → E95_S4 | Loki | Diffusion_1.0 | 0.1803 | 0.8584 | 0.3481 |
| E95_S3 → E95_S4 | Loki | Diffusion_0.1 | 0.2573 | 0.8775 | 0.3572 |
| E95_S3 → E95_S4 | Loki | Diffusion_0.01 | 0.2932 | 0.8855 | 0.3601 |
| E95_S3 → E95_S4 | Loki | Diffusion_0.001 | 0.3201 | 0.8857 | 0.3625 |
| E95_S3 → E95_S4 | Loki | Diffusion_0.00001 | 0.3205 | 0.8864 | 0.3622 |

Table S9: Nicheformer's performance on mouse embryonic reconstruction

| Experiment | Foundation model | Method | LTARI | Pearson | Kendall |
|---|---|---|---|---|---|
| E95_S1 → E95_S2 | Nicheformer | Original | 0.2312 | 0.8785 | 0.359 |
| E95_S1 → E95_S2 | Nicheformer | Diffusion_5.0 | 0.1016 | 0.8518 | 0.3396 |
| E95_S1 → E95_S2 | Nicheformer | Diffusion_2.0 | 0.0992 | 0.8458 | 0.339 |
| E95_S1 → E95_S2 | Nicheformer | Diffusion_100.0 | 0.073 | 0.838 | 0.3264 |
| E95_S1 → E95_S2 | Nicheformer | Diffusion_10.0 | 0.0991 | 0.8514 | 0.3395 |
| E95_S1 → E95_S2 | Nicheformer | Diffusion_1.0 | 0.0962 | 0.8463 | 0.3395 |
| E95_S1 → E95_S2 | Nicheformer | Diffusion_0.1 | 0.1009 | 0.8512 | 0.3377 |
| E95_S1 → E95_S2 | Nicheformer | Diffusion_0.01 | 0.0999 | 0.85 | 0.3387 |
| E95_S1 → E95_S2 | Nicheformer | Diffusion_0.001 | 0.0877 | 0.8396 | 0.3356 |
| E95_S1 → E95_S2 | Nicheformer | Diffusion_0.0001 | 0.0829 | 0.8497 | 0.3429 |
| E95_S1 → E95_S2 | Nicheformer | Diffusion_0.00001 | 0.1049 | 0.8526 | 0.3402 |
| E95_S2 → E95_S3 | Nicheformer | Original | 0.1703 | 0.8497 | 0.3414 |
| E95_S2 → E95_S3 | Nicheformer | Diffusion_5.0 | 0.1649 | 0.8412 | 0.3355 |
| E95_S2 → E95_S3 | Nicheformer | Diffusion_2.0 | 0.1622 | 0.8438 | 0.3376 |
| E95_S2 → E95_S3 | Nicheformer | Diffusion_100.0 | 0.1956 | 0.8357 | 0.3394 |
| E95_S2 → E95_S3 | Nicheformer | Diffusion_10.0 | 0.1656 | 0.8386 | 0.3352 |
| E95_S2 → E95_S3 | Nicheformer | Diffusion_1.0 | 0.1622 | 0.8447 | 0.339 |
| E95_S2 → E95_S3 | Nicheformer | Diffusion_0.1 | 0.1616 | 0.8453 | 0.3389 |
| E95_S2 → E95_S3 | Nicheformer | Diffusion_0.01 | 0.173 | 0.8431 | 0.3388 |
| E95_S2 → E95_S3 | Nicheformer | Diffusion_0.001 | 0.2226 | 0.8259 | 0.3322 |
| E95_S2 → E95_S3 | Nicheformer | Diffusion_0.0001 | 0.1085 | 0.8117 | 0.3191 |
| E95_S2 → E95_S3 | Nicheformer | Diffusion_0.00001 | 0.1655 | 0.849 | 0.3401 |
| E95_S3 → E95_S4 | Nicheformer | Original | 0.0939 | 0.7857 | 0.3255 |
| E95_S3 → E95_S4 | Nicheformer | Diffusion_5.0 | 0.3051 | 0.8896 | 0.3621 |
| E95_S3 → E95_S4 | Nicheformer | Diffusion_2.0 | 0.2823 | 0.8823 | 0.3591 |
| E95_S3 → E95_S4 | Nicheformer | Diffusion_100.0 | 0.1828 | 0.8686 | 0.3485 |
| E95_S3 → E95_S4 | Nicheformer | Diffusion_10.0 | 0.2903 | 0.8851 | 0.3605 |
| E95_S3 → E95_S4 | Nicheformer | Diffusion_1.0 | 0.2384 | 0.8728 | 0.3568 |
| E95_S3 → E95_S4 | Nicheformer | Diffusion_0.1 | 0.2545 | 0.8767 | 0.357 |
| E95_S3 → E95_S4 | Nicheformer | Diffusion_0.01 | 0.2338 | 0.8721 | 0.3561 |
| E95_S3 → E95_S4 | Nicheformer | Diffusion_0.001 | 0.2487 | 0.8749 | 0.3562 |
| E95_S3 → E95_S4 | Nicheformer | Diffusion_0.0001 | 0.1183 | 0.7814 | 0.326 |
| E95_S3 → E95_S4 | Nicheformer | Diffusion_0.00001 | 0.1031 | 0.7656 | 0.3213 |

### S11 Few-Shot Clustering Results

Table S10 illustrates the dataset-specific few-shot clustering performance of scGPT and scFoundation and their DRIFT-enhanced versions after fine-tuning on only one train dataset. For few-shot learning, we perform a task similar to GraphST, where the model aims to reconstruct the input gene-expression profile. DRIFT-incorporated scFoundation's mean ARI performance shows almost no improvement from 0.523 in the zero-shot setting to 0.525 in the few-shot setting. However, DRIFT-incorporated scGPT displays major improvement with fine-tuning, with its mean ARI improving from 0.447 in the zero-shot setting to 0.5556 in the few-shot setting.

Table S10: **Clustering Results** - The table lists the mean few-shot ARI performance of scGPT, scFoundation, and their DRIFT-incorporated versions on every dataset. These results show that few-shot training can improve the performance of some foundation models.

| | Original scGPT | DRIFT scGPT | Original scFoundation | DRIFT scFoundation |
|---|---|---|---|---|
| 10xHPC | 0.3193 | 0.5904 | 0.3103 | 0.4829 |
| MERHH | 0.6207 | 0.5779 | 0.4244 | 0.5358 |
| MERMPH | 0.2967 | 0.5076 | 0.1712 | 0.5190 |
| MERMB | 0.4301 | 0.4118 | 0.3582 | 0.4858 |
| MERMBA | 0.3511 | 0.6901 | 0.3972 | 0.6014 |

### Figures

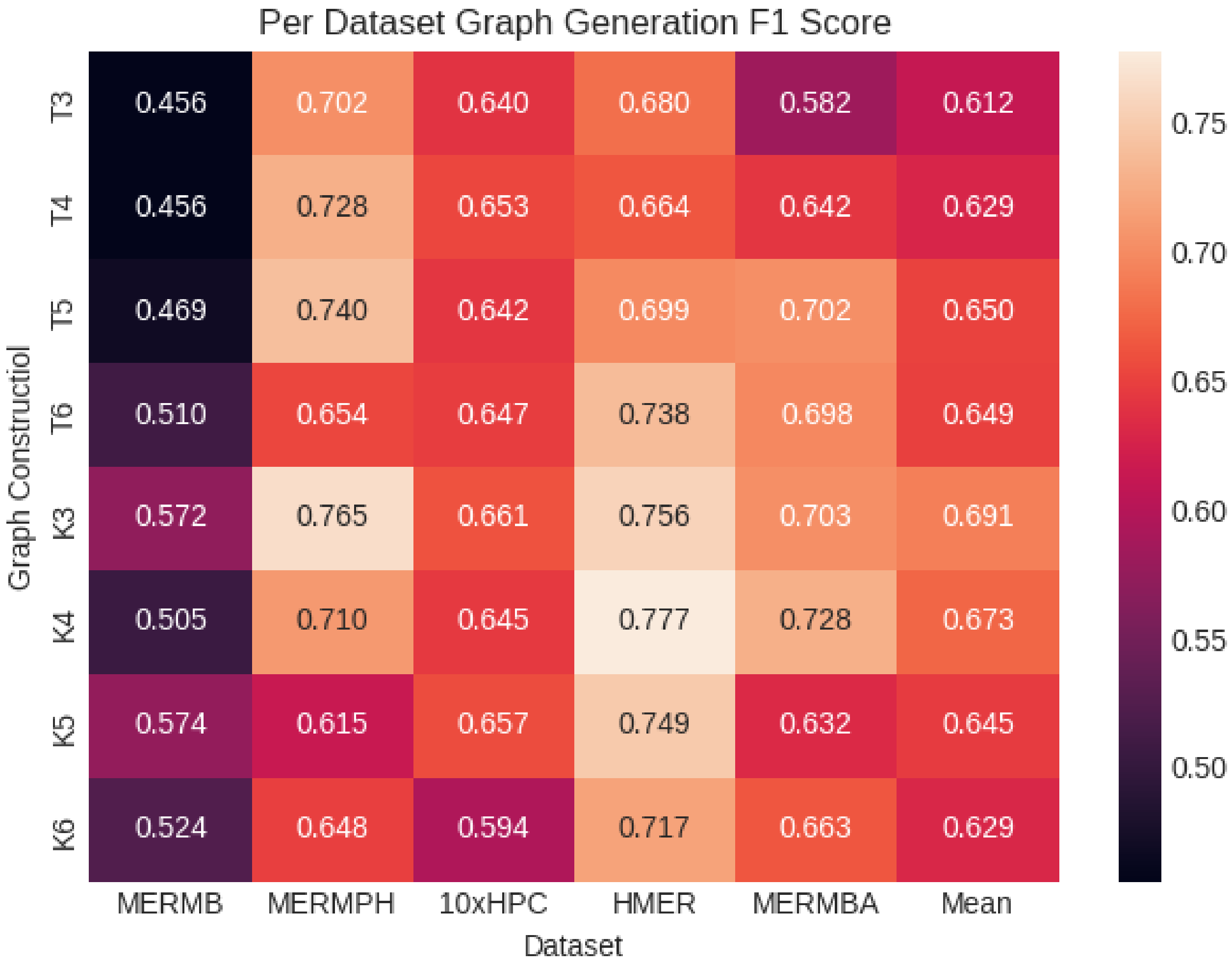


Figure S1: F1-scores of different graph construction techniques for scGPT over all datasets in the cell annotation task.

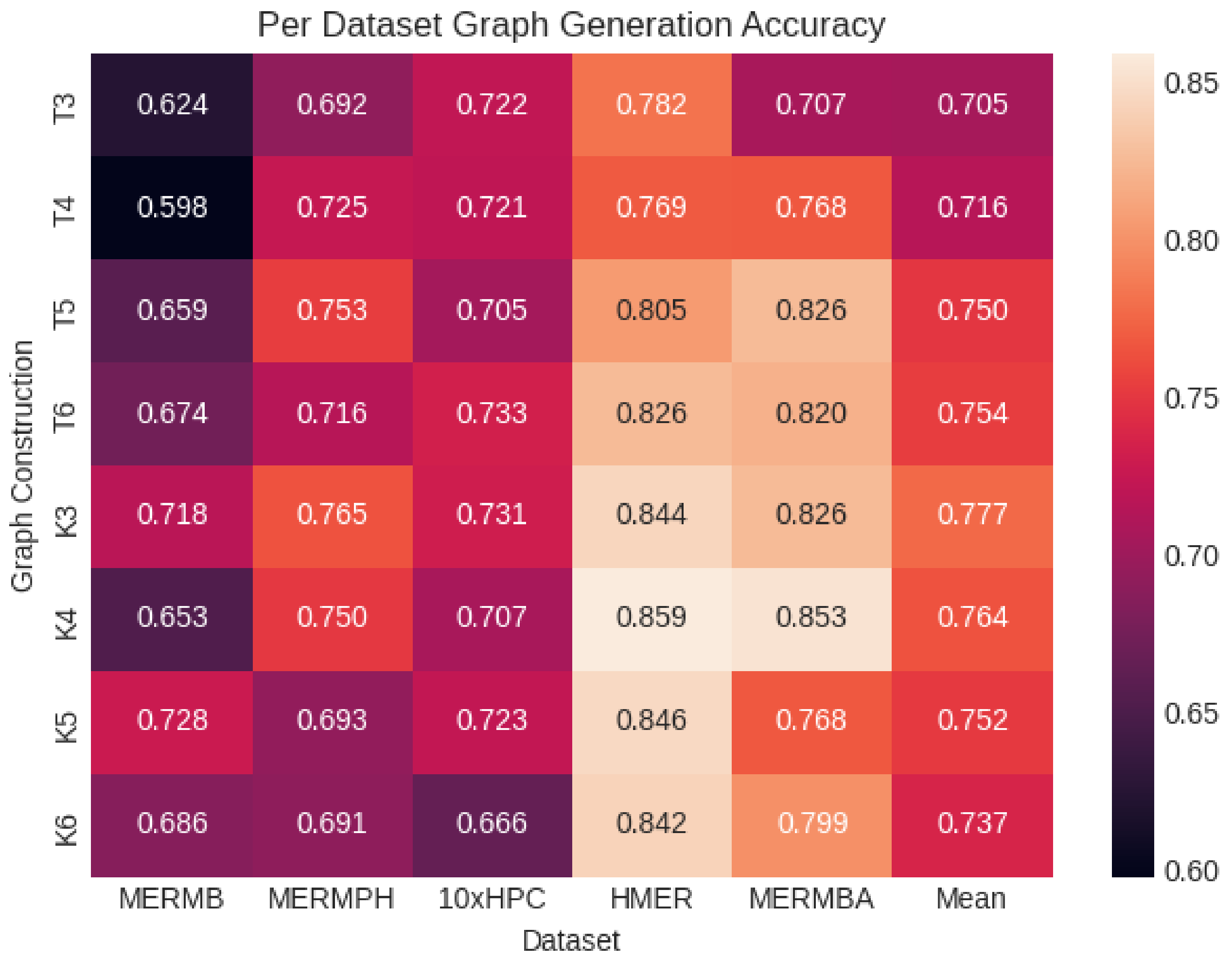


Figure S2: Accuracy of different graph construction techniques for scGPT over all datasets in the cell annotation task.

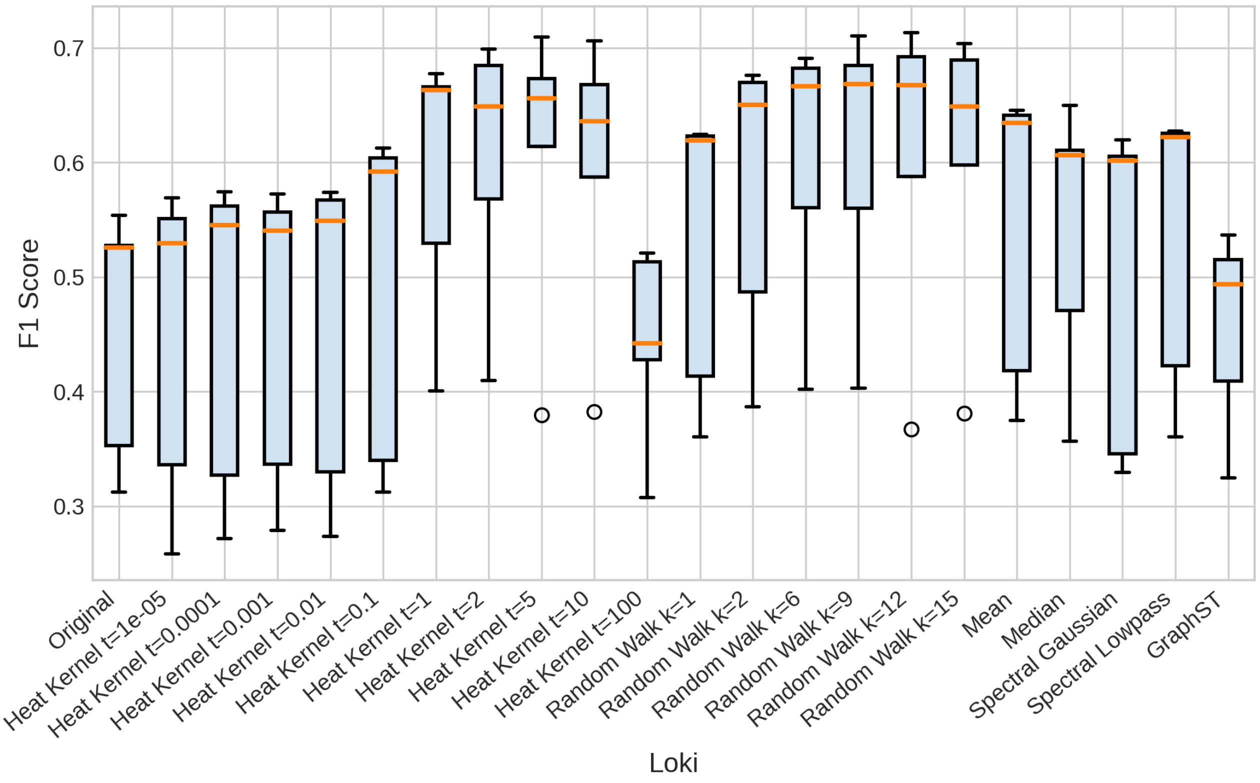


Figure S3: F1-scores of different diffusion methods for Loki over all datasets in the cell annotation task.

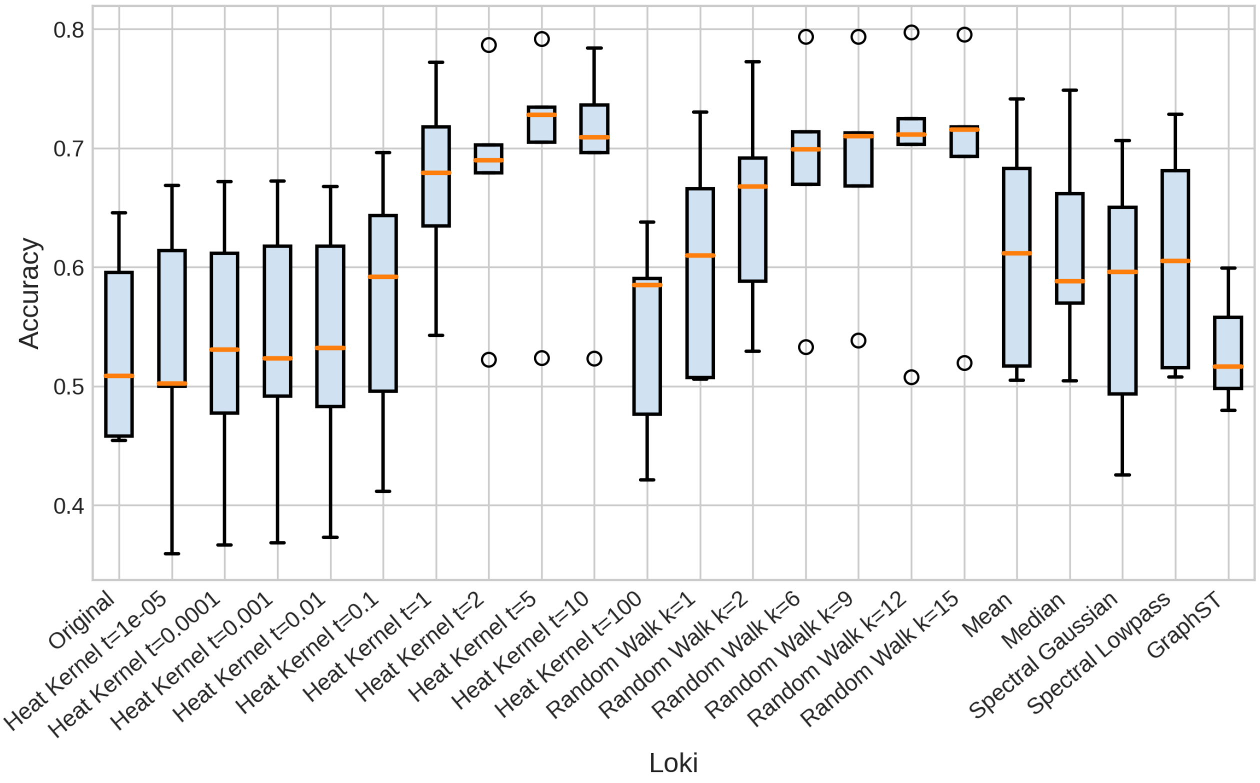


Figure S4: Accuracy of different diffusion methods for Loki over all datasets in the cell annotation task.

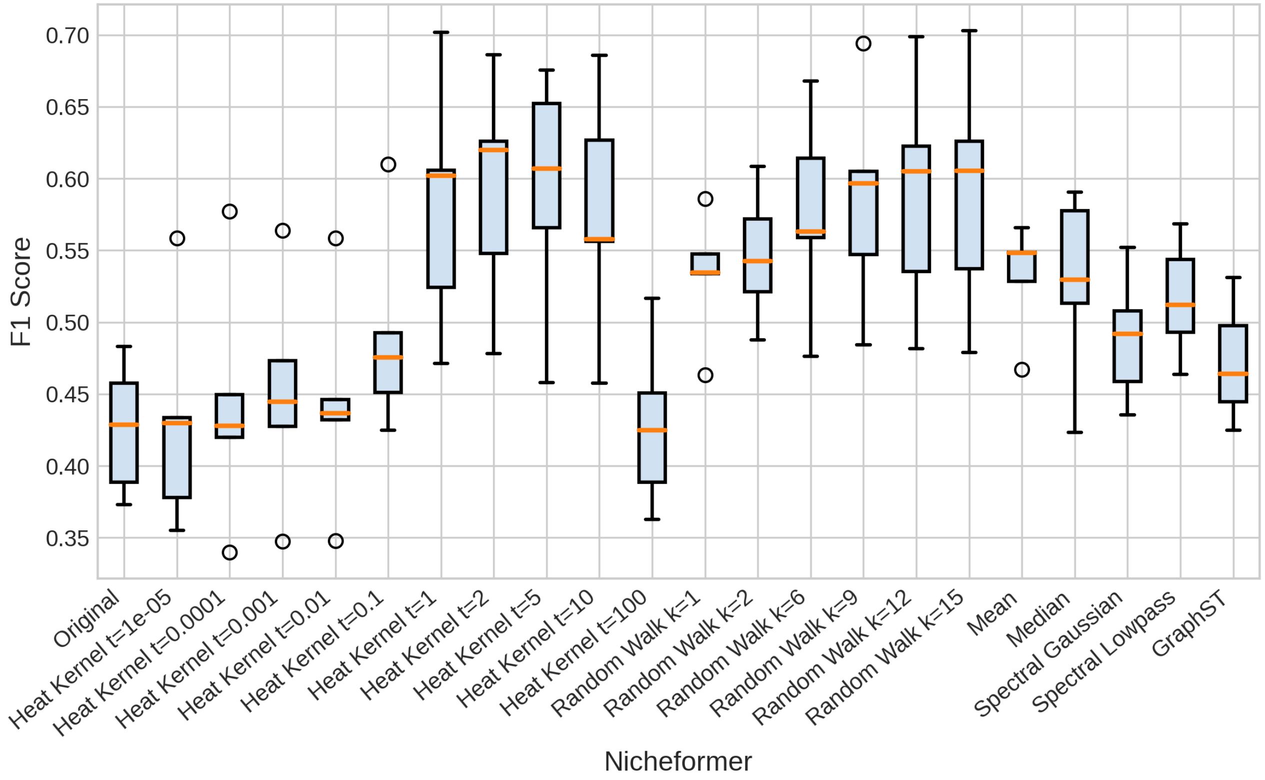


Figure S5: F1-scores of different diffusion methods for Nicheformer over all datasets in the cell annotation task.

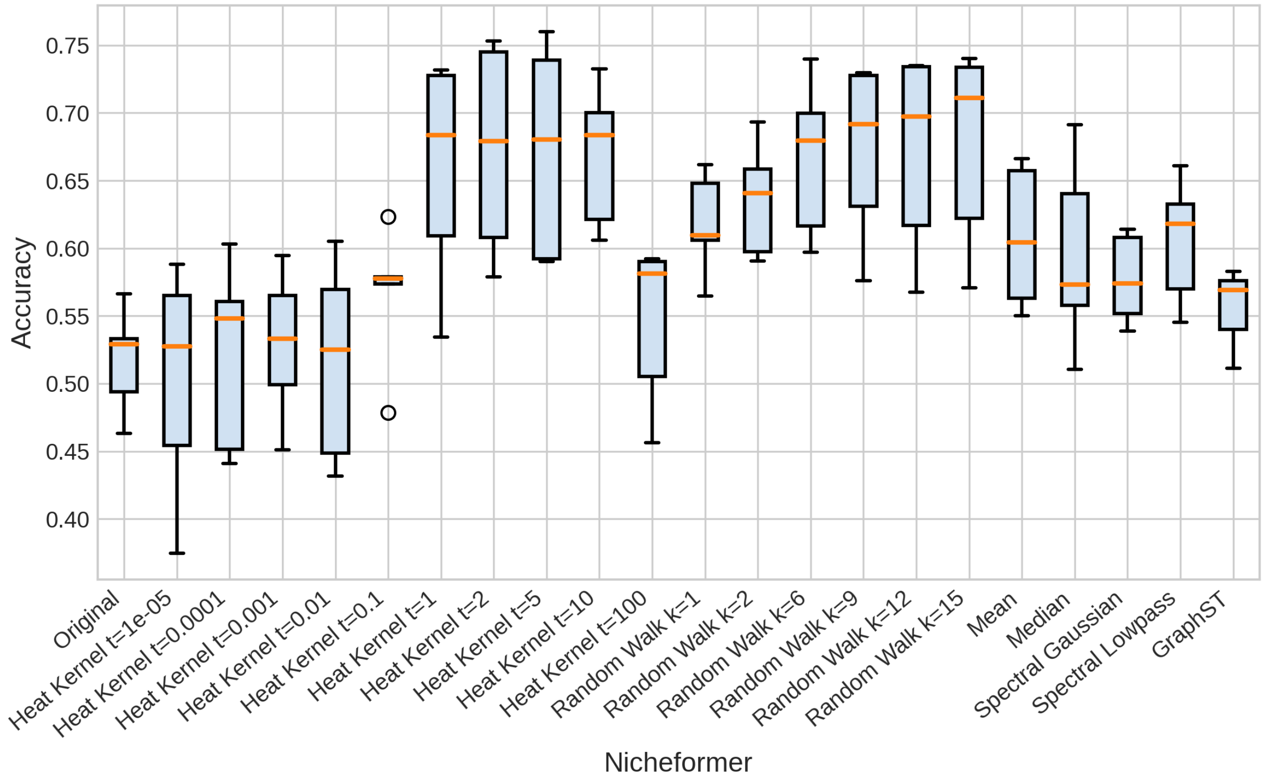


Figure S6: Accuracy of different diffusion methods for Nicheformer over all datasets in the cell annotation task.

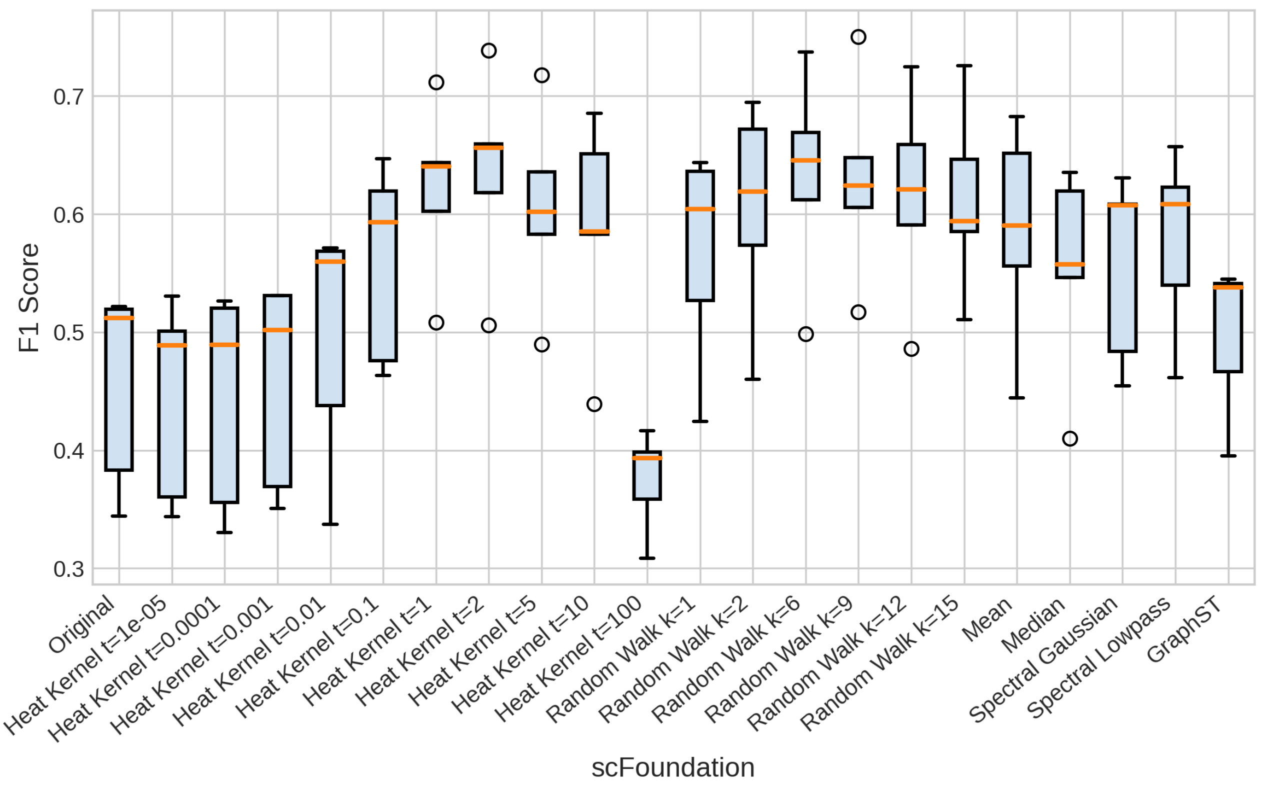


Figure S7: F1-scores of different diffusion methods for scFoundation over all datasets in the cell annotation task.

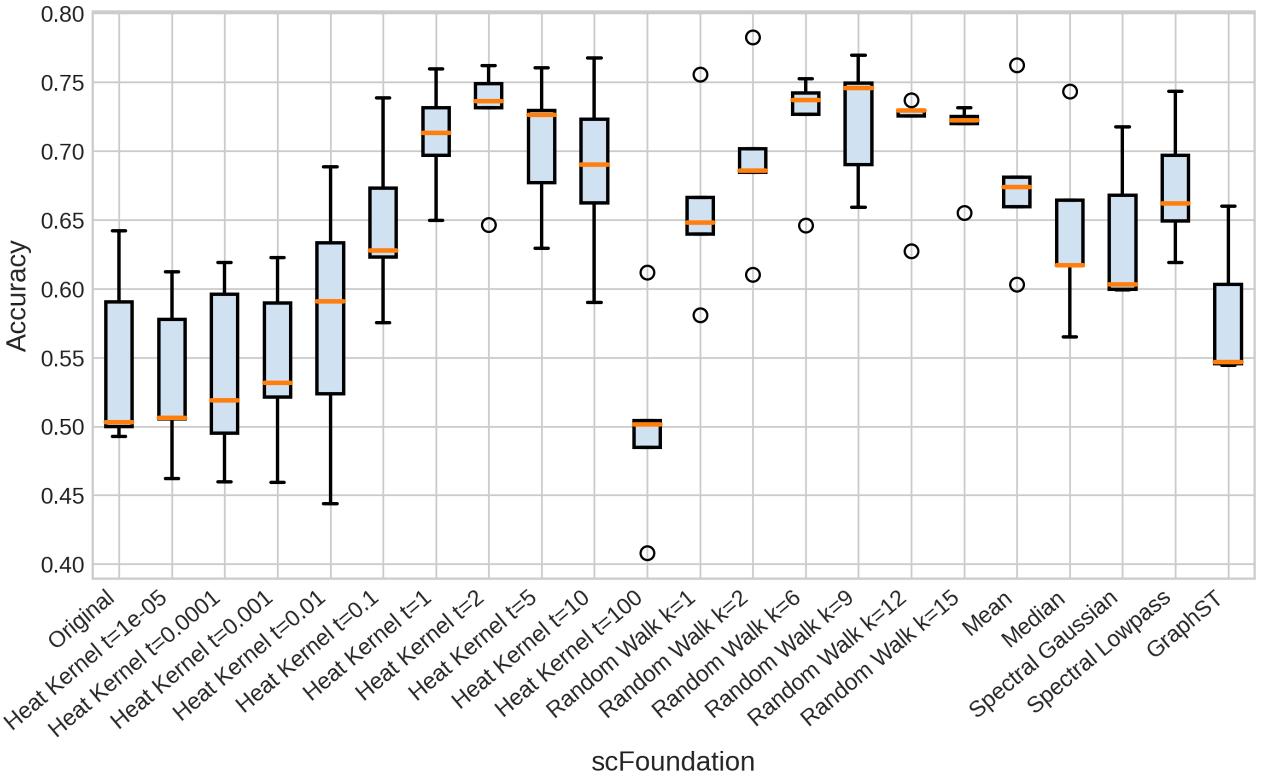


Figure S8: Accuracy of different diffusion methods for scFoundation over all datasets in the cell annotation task.

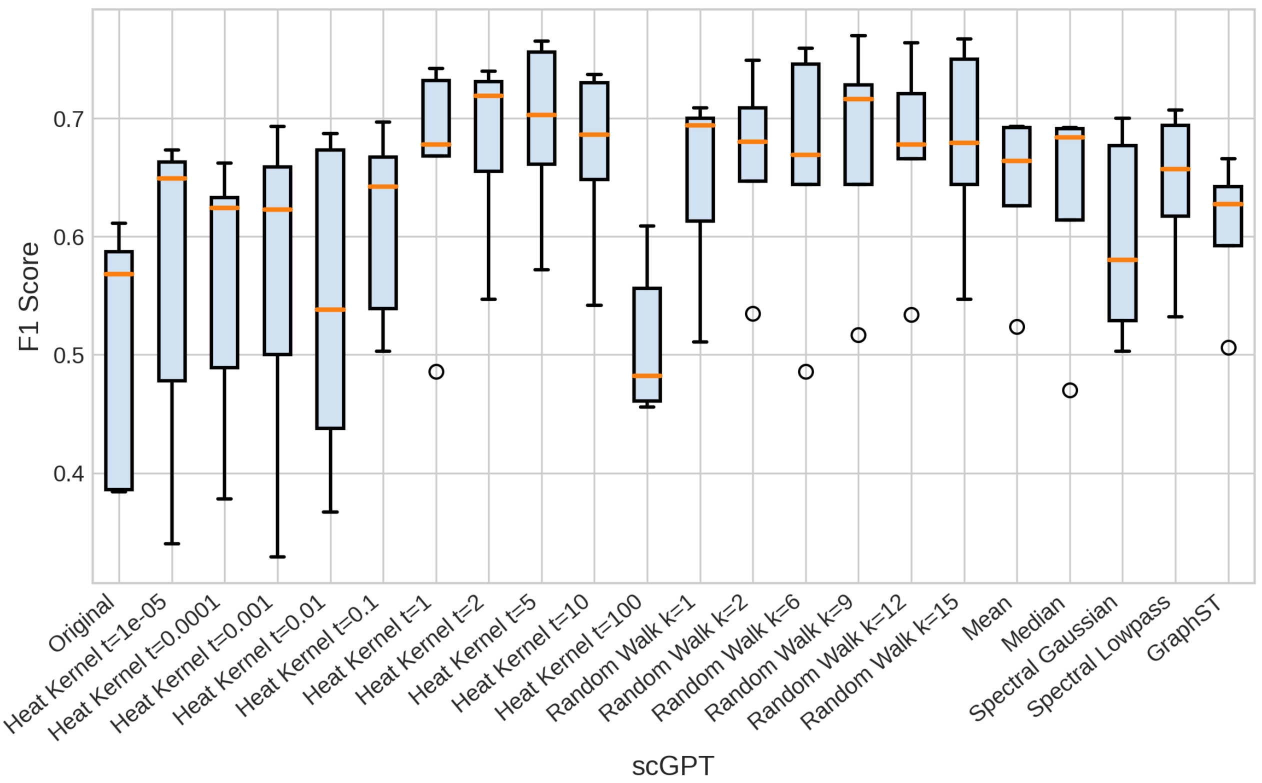


Figure S9: F1-scores of different diffusion methods for scGPT over all datasets in the cell annotation task.

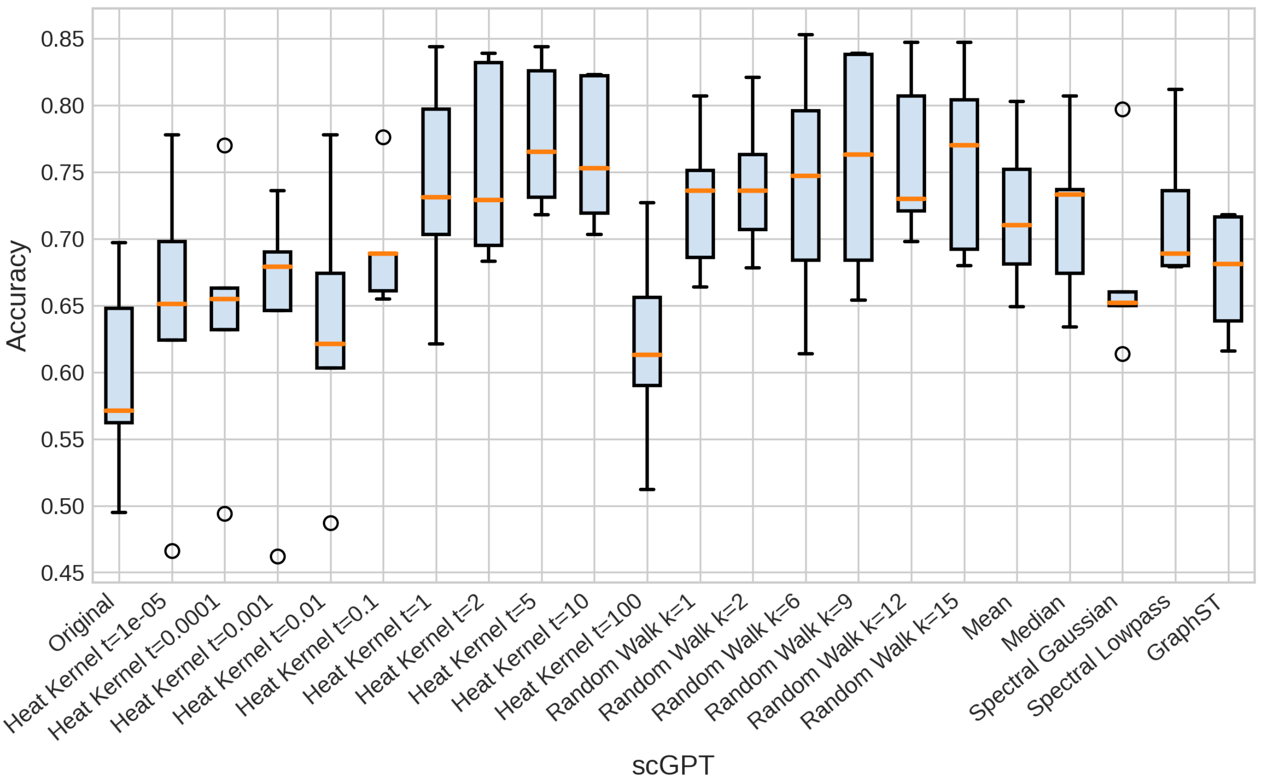


Figure S10: Accuracy of different diffusion methods for scGPT over all datasets in the cell annotation task.

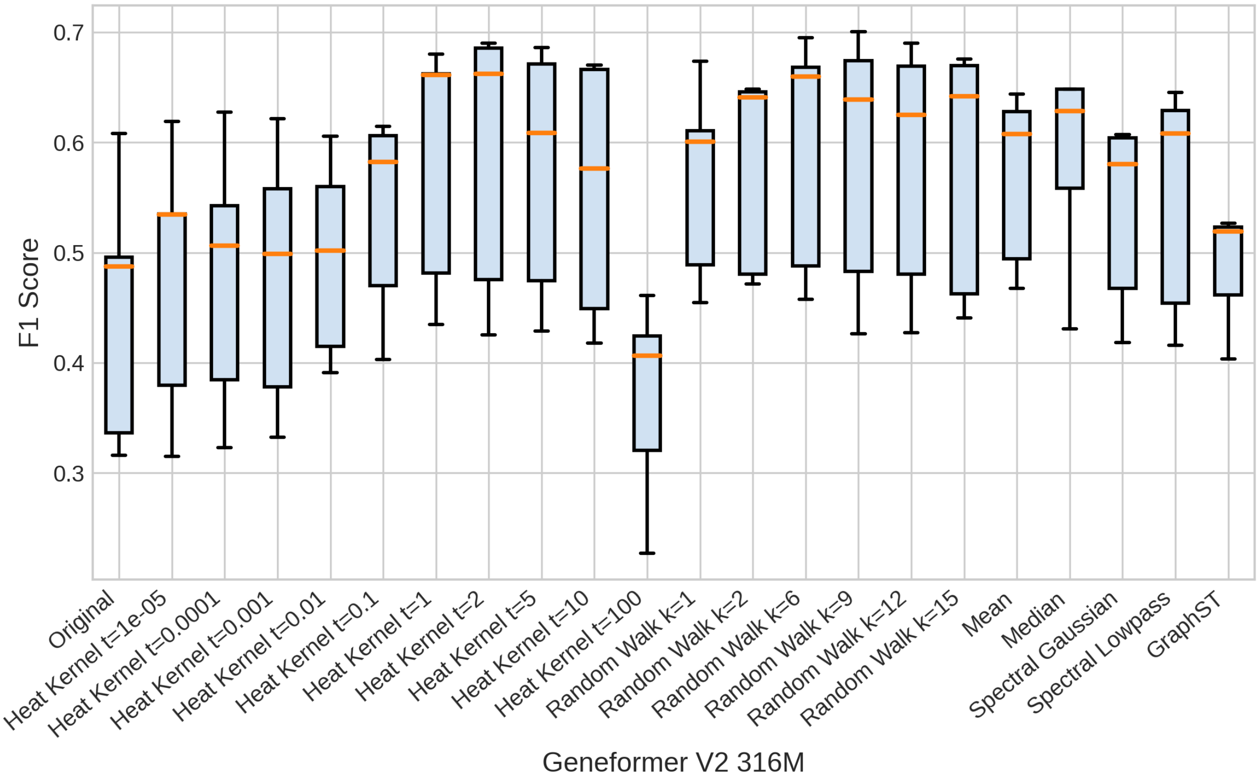


Figure S11: F1-scores of different diffusion methods for Geneformer V2 with 316 million parameters over all datasets in the cell annotation task.

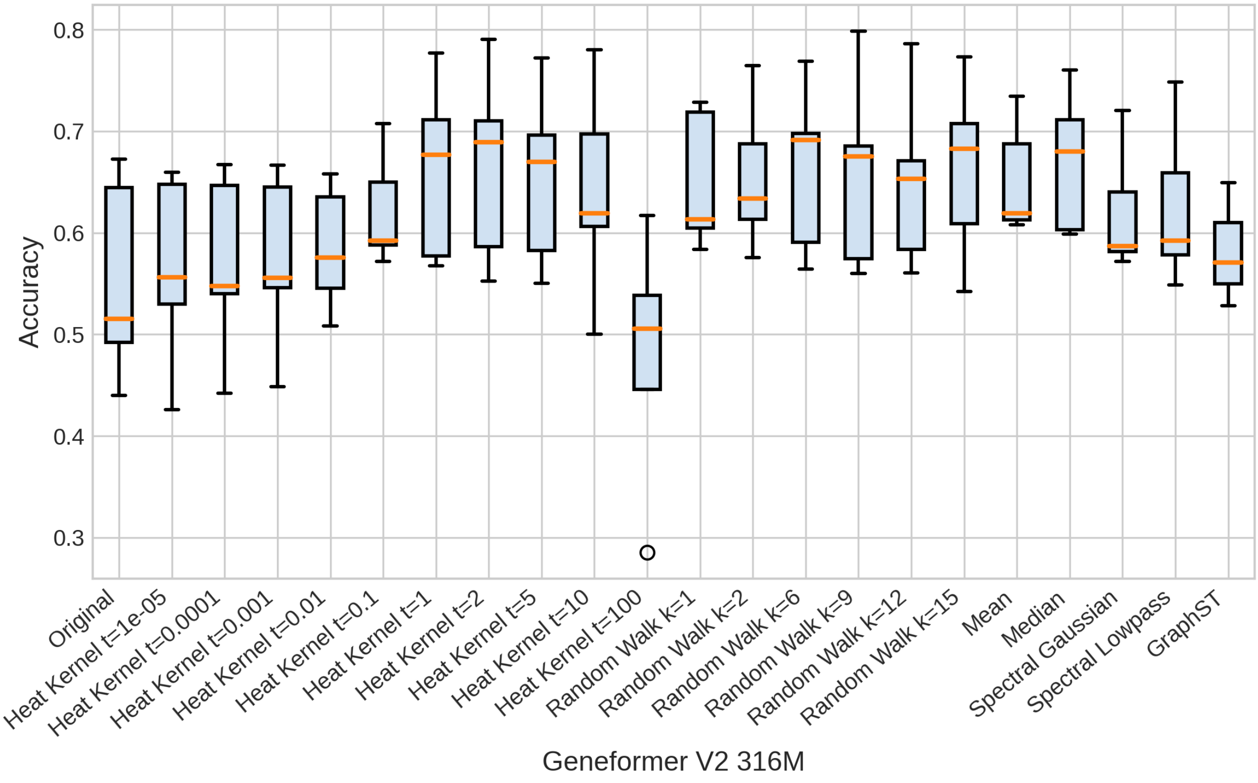


Figure S12: Accuracy of different diffusion methods for Geneformer V2 with 316 million parameters over all datasets in the cell annotation task.

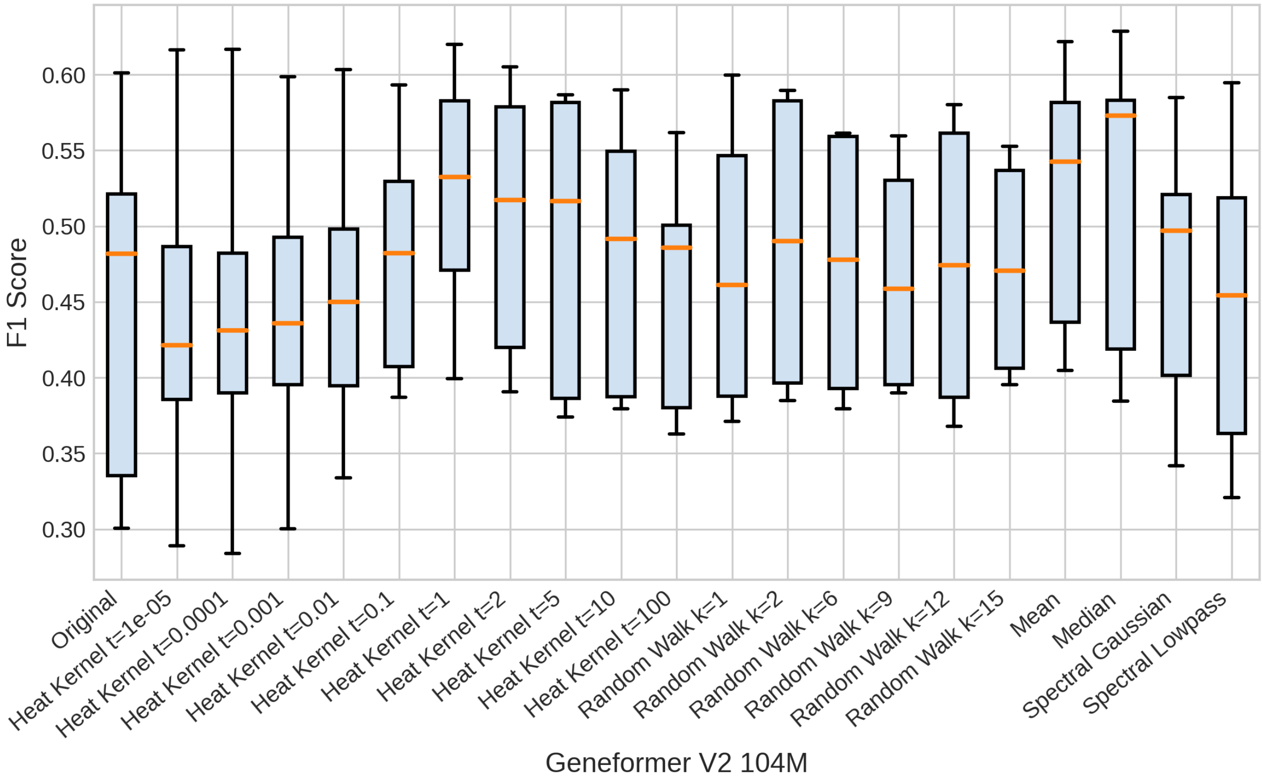


Figure S13: F1-scores of different diffusion methods for Geneformer V2 with 104 million parameters over all datasets in the cell annotation task.

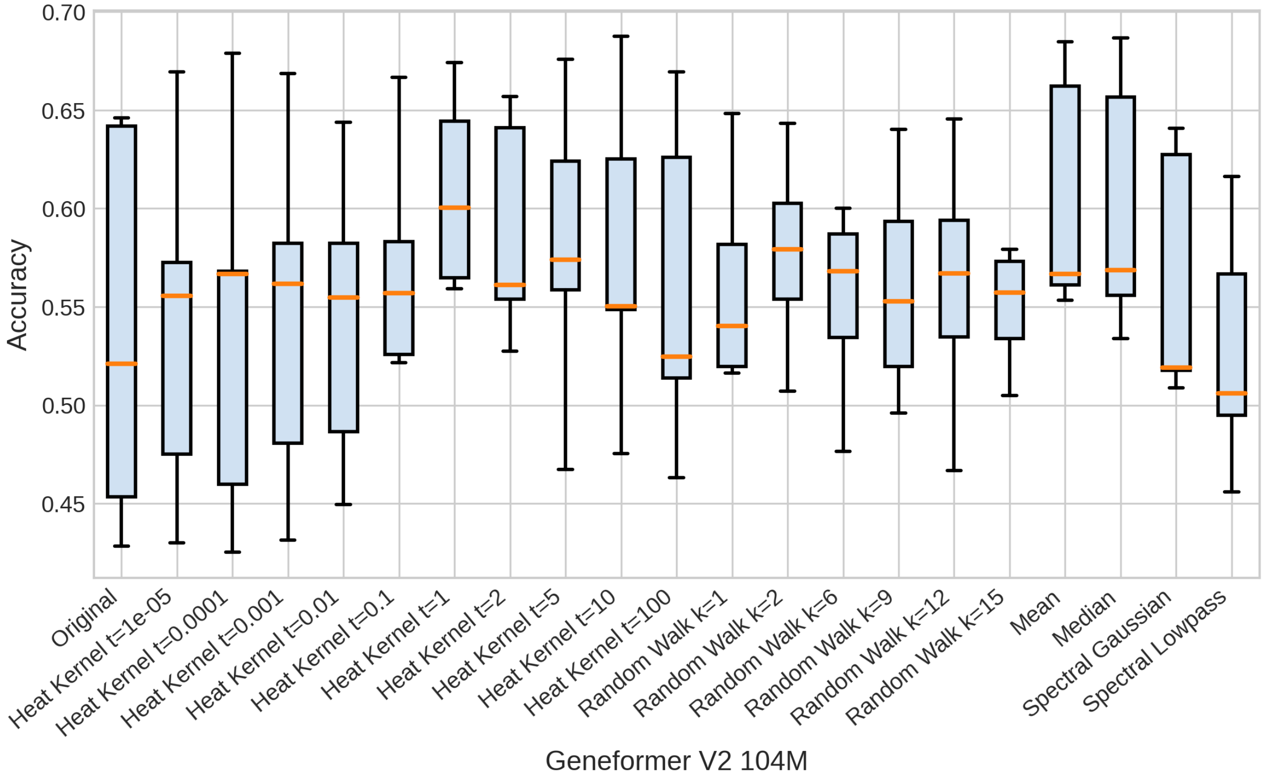


Figure S14: Accuracy of different diffusion methods for Geneformer V2 with 104 million parameters over all datasets in the cell annotation task.

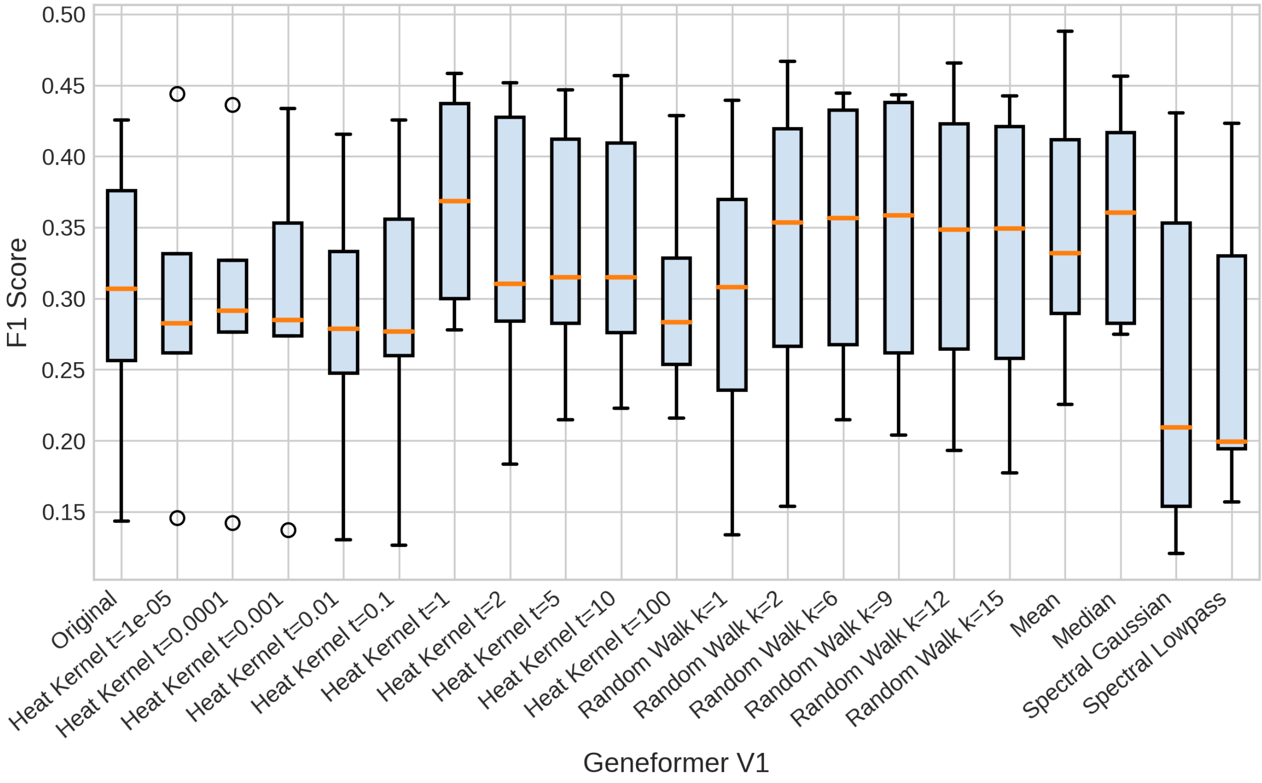


Figure S15: F1-scores of different diffusion methods for Geneformer V1 over all datasets in the cell annotation task.

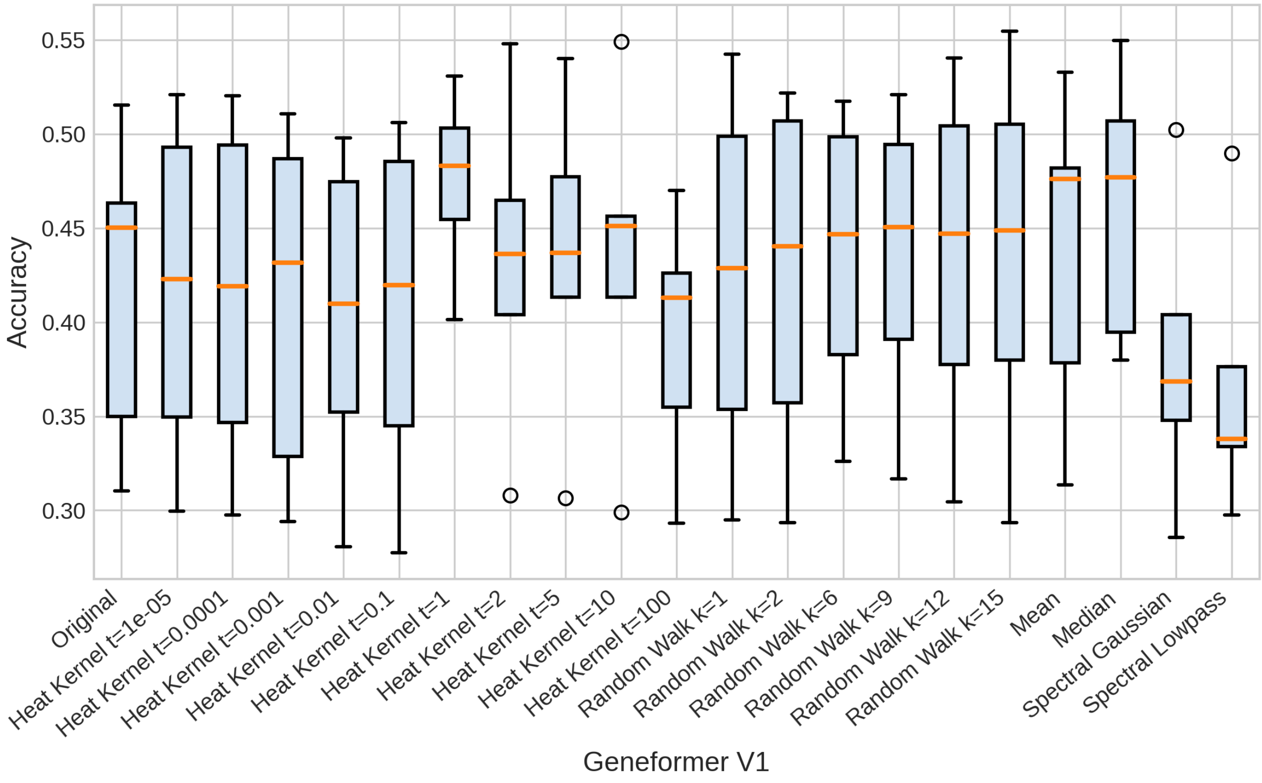


Figure S16: Accuracy of different diffusion methods for Geneformer V1 over all datasets in the cell annotation task.

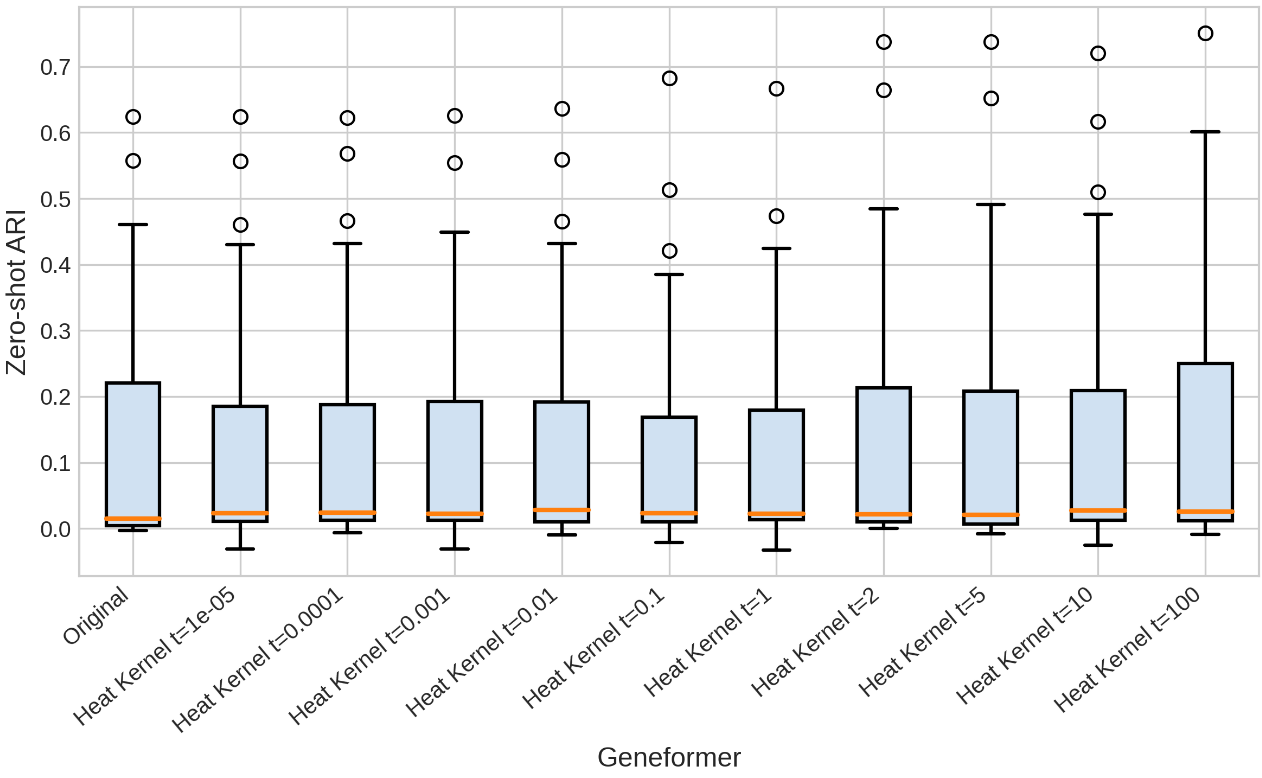


Figure S17: Zero-shot clustering ARI of different values of $t$ for Geneformer over all datasets.

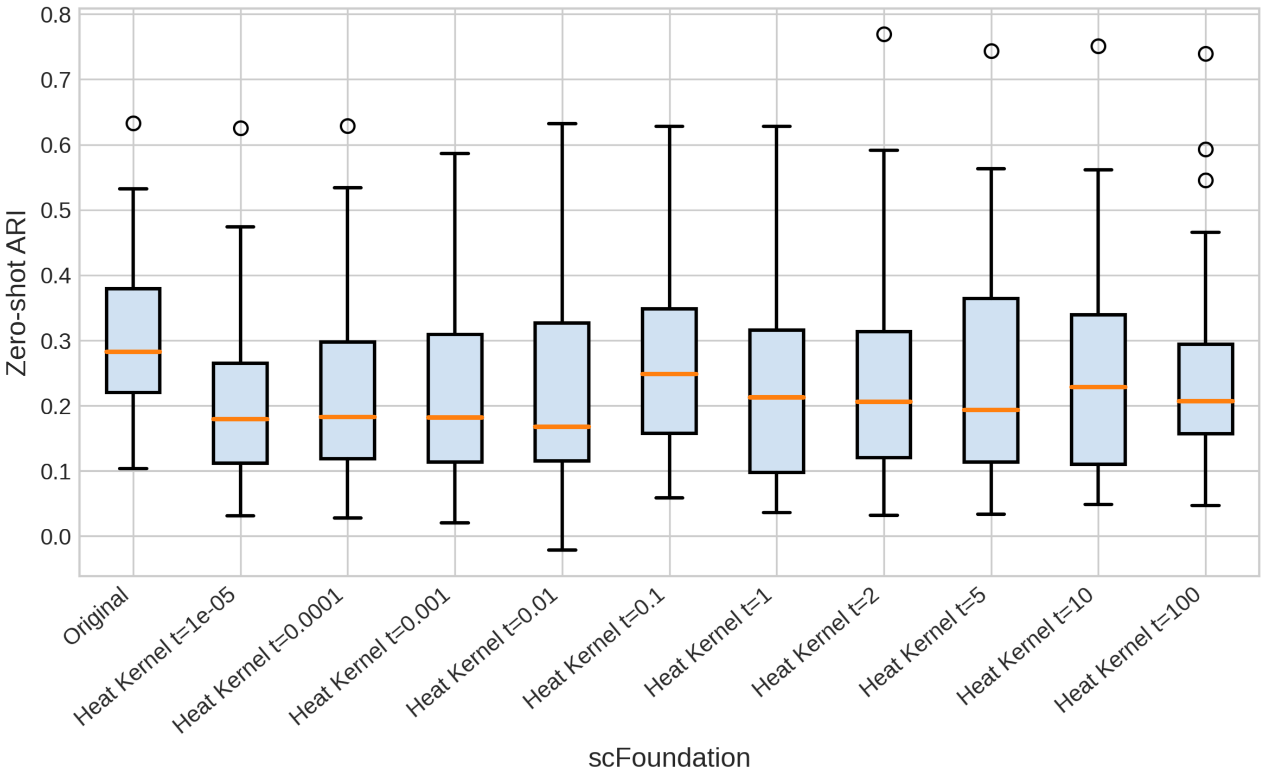


Figure S18: Zero-shot clustering ARI of different values of $t$ for scFoundation over all datasets.

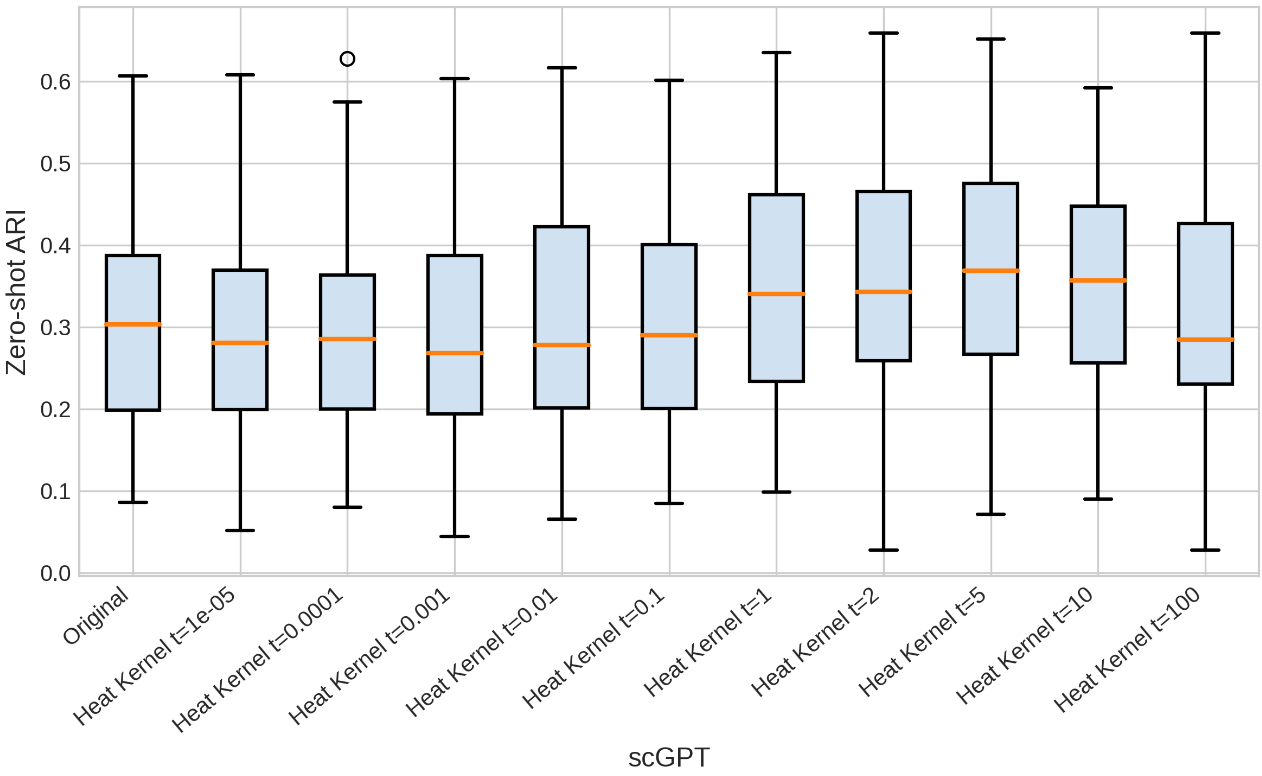


Figure S19: Zero-shot clustering ARI of different values of $t$ for scGPT over all datasets.

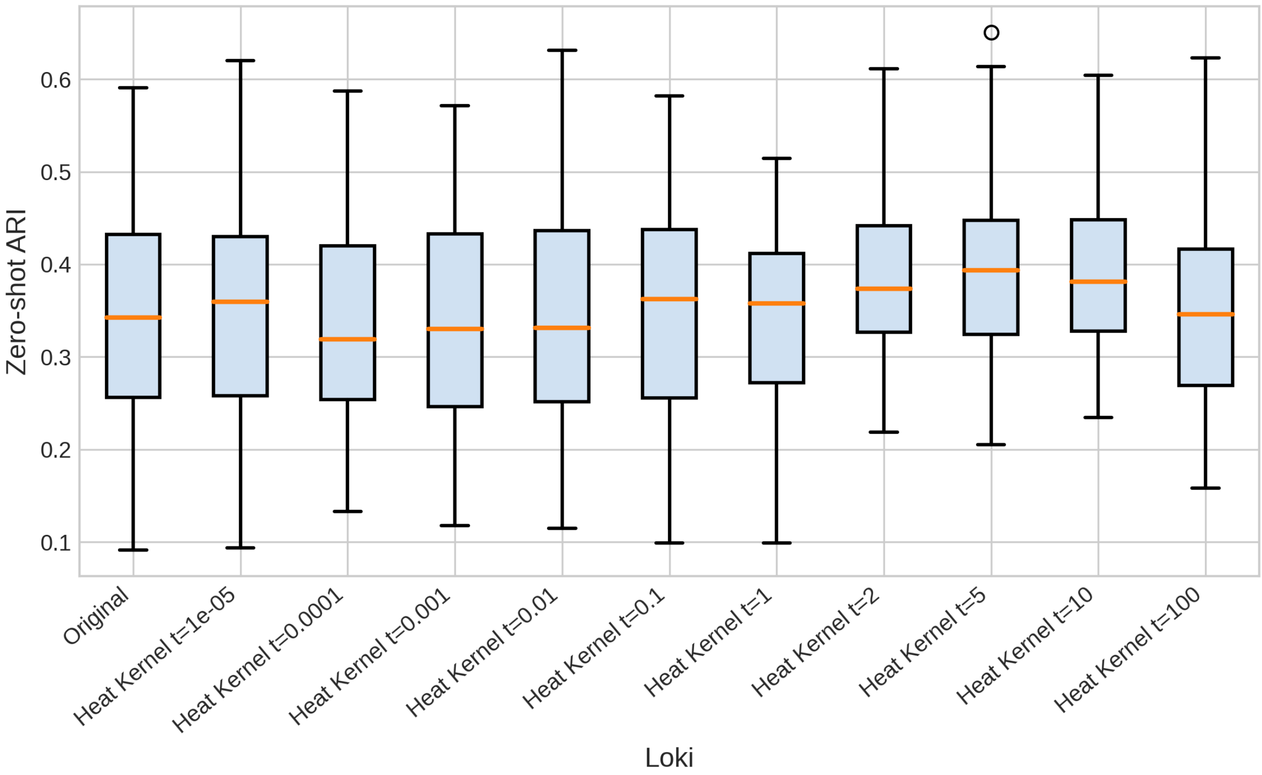


Figure S20: Zero-shot clustering ARI of different values of $t$ for Loki over all datasets.

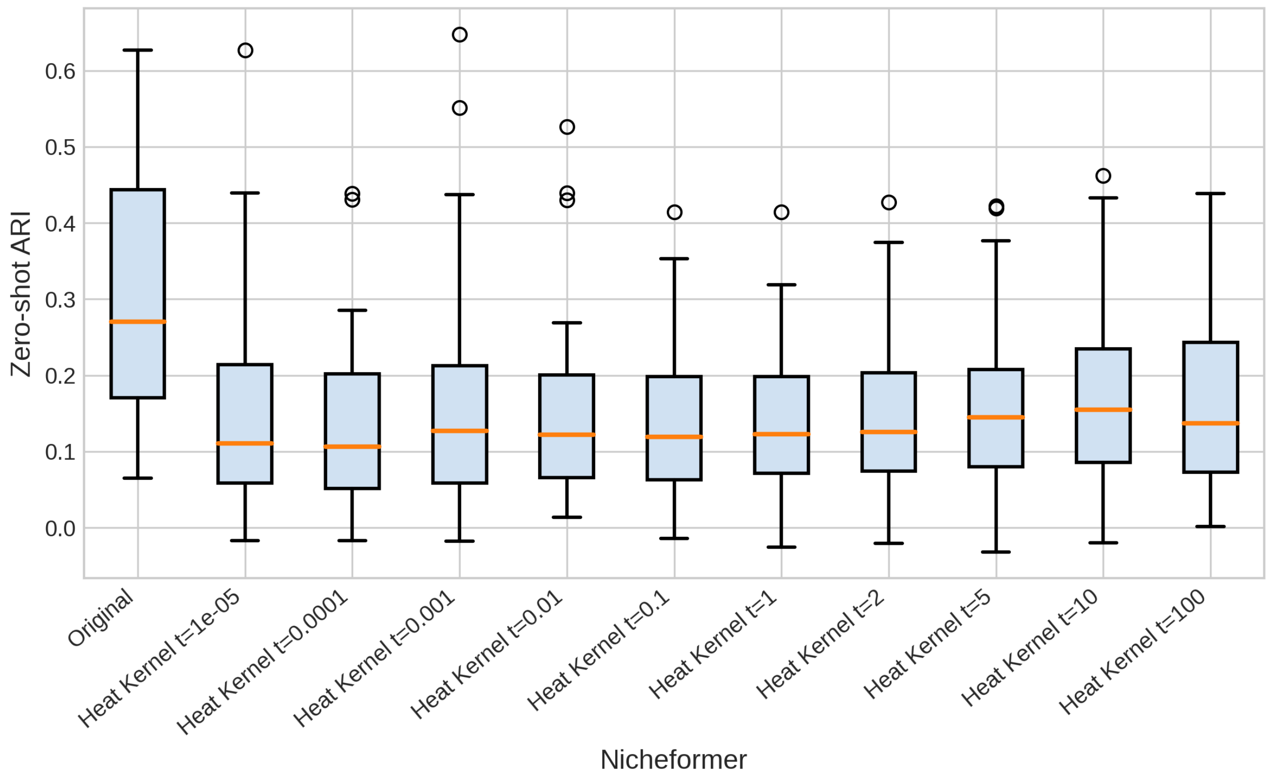


Figure S21: Zero-shot clustering ARI of different values of $t$ for Nicheformer over all datasets.

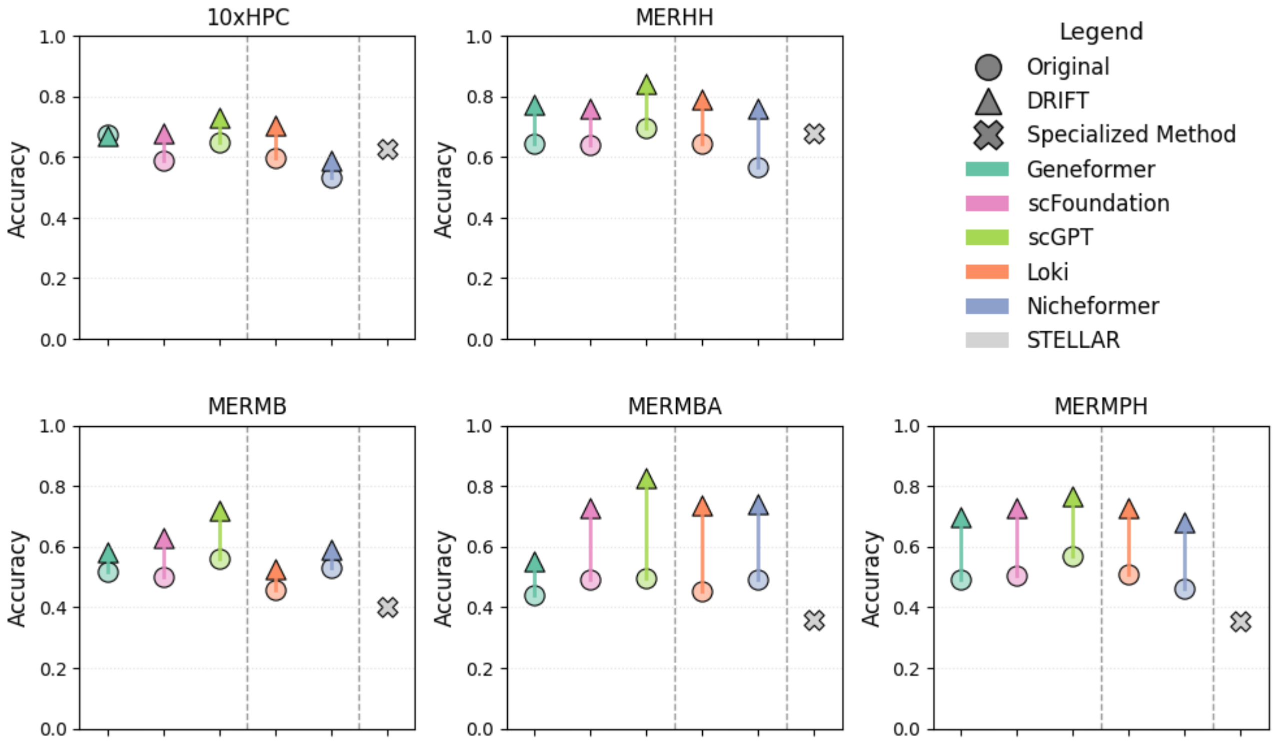


Figure S22: **Annotation Results** - A quantitative comparison across multiple datasets. The circles represent the performance of the original foundation models, the triangles represent the DRIFT-incorporated foundation models, and the X represents the specialized method STELLAR. The colors specify which column corresponds to which foundation model (the order of models in the legend is consistent with column placement). This figure demonstrates that DRIFT improves the F1-score performance of foundation models on the annotation task. Furthermore, DRIFT-incorporated foundation models, specifically scGPT show state-of-the-art performance, surpassing even the specialized model - STELLAR.

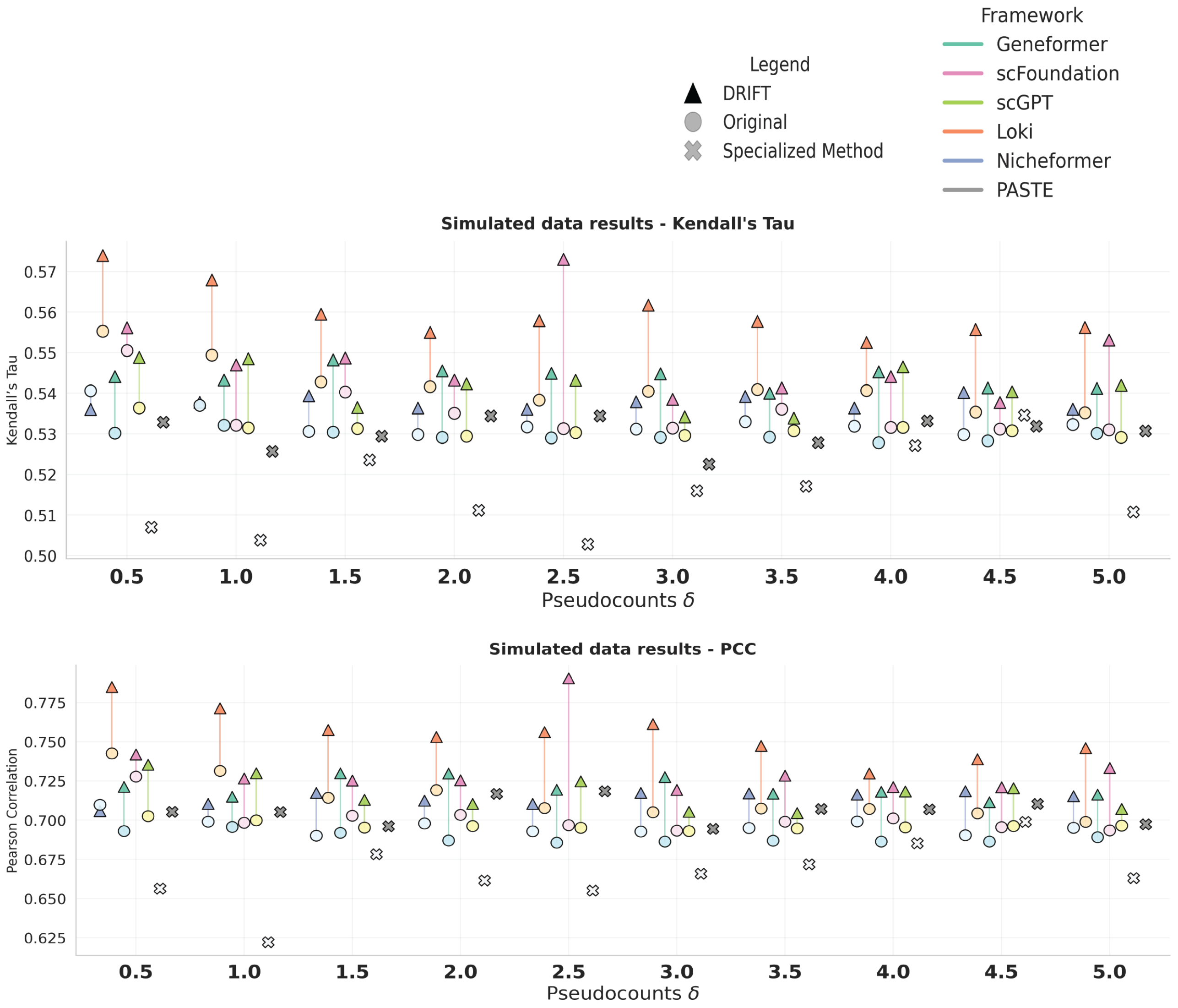


Figure S23: **Simulated Data Alignment Results** - Our results show that with DRIFT-enhanced embeddings, alignments consistently show higher PCC and Kendall's $\tau$ values, indicating better alignments. This is also consistent with the Translation Error metric in Section 3.2 of the main text.

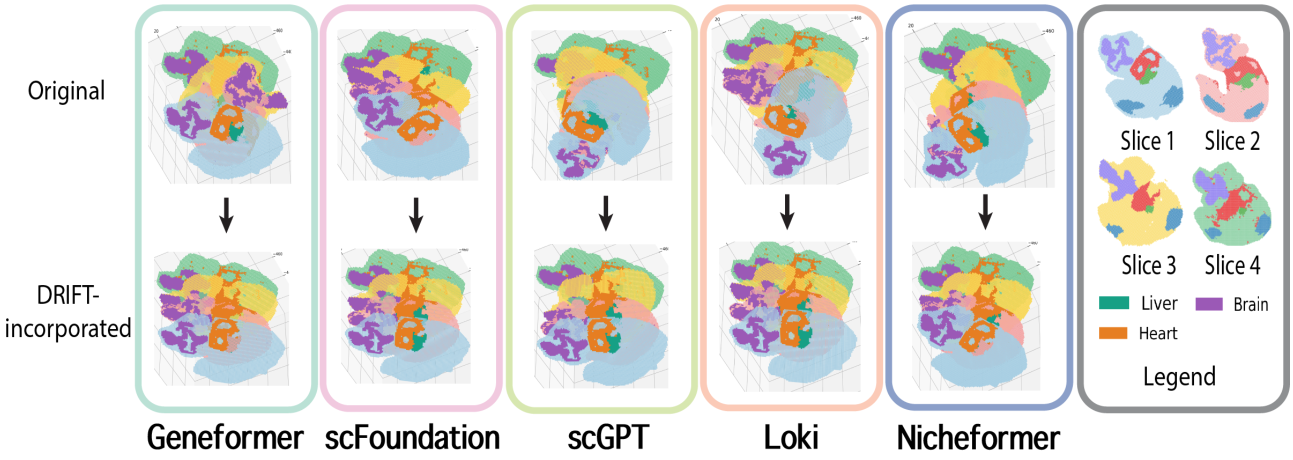


Figure S24: 3D reconstruction from all foundation models with and without DRIFT
